# CDK phosphorylation of *Xenopus laevis* M18BP1 promotes its metaphase centromere localization

**DOI:** 10.1101/355487

**Authors:** Bradley T. French, Aaron F. Straight

## Abstract

Chromosome segregation requires the centromere, the site on chromosomes where kinetochores assemble in mitosis to attach chromosomes to the mitotic spindle. Centromere identity is defined epigenetically by the presence of nucleosomes containing the histone H3 variant CENP-A. New CENP-A nucleosome assembly occurs at the centromere every cell cycle during G1, but how CENP-A nucleosome assembly is spatially and temporally restricted remains poorly understood. Centromere recruitment of factors required for CENP-A assembly is mediated in part by the three-protein Mis18 complex (Mis18α, Mis18β, M18BP1). Here we show that *Xenopus* M18BP1 localizes to centromeres during metaphase - prior to CENP-A assembly - by binding to CENP-C using a highly conserved SANTA domain. We find that Cdk phosphorylation of M18BP1 is necessary for M18BP1 to bind CENP-C and localize to centromeres in metaphase. Surprisingly, mutations which disrupt the metaphase M18BP1/CENP-C interaction cause defective nuclear localization of M18BP1 in interphase, resulting in defective CENP-A nucleosome assembly. We propose that M18BP1 may identify centromeric sites in metaphase for subsequent CENP-A nucleosome assembly in interphase.

## Introduction

During cell division, eukaryotes segregate their genomes by attaching chromosomes to the microtubules of the mitotic spindle through the kinetochore. Kinetochores assemble in mitosis on a specialized region of the chromosome termed the centromere. Mutations that disrupt the functions of centromere and kinetochore proteins cause chromosome missegregation, genome instability, and cell death. Centromeres are distinguished by the incorporation of a histone H3 variant termed CENP-A (CENtromere Protein A) into centromeric nucleosomes. CENP-A nucleosomes epigenetically determine the function of centromeres, and loss of CENP-A results in defective kinetochore assembly and improper chromosome segregation. Thus, a central question in chromosome segregation is how cells selectively maintain CENP-A chromatin at the centromere.

CENP-A nucleosomes are equally distributed to each sister chromatid during DNA replication (Jansen et al., 2007). To prevent the replication-coupled dilution of CENP-A chromatin, new CENP-A nucleosomes are assembled once per cell cycle during G1. New CENP-A assembly is mediated by HJURP (Holliday junction-recognizing protein), the histone chaperone that binds soluble CENP-A/H4 dimers (Dunleavy et al., 2009; Foltz et al., 2009). HJURP localization is sufficient to promote local CENP-A deposition so its localization must be restricted to centromeres (Barnhart et al., 2011; French et al., 2017; Shono et al., 2015). Centromere-specific HJURP targeting requires a CENP-A nucleosome-binding protein, CENP-C (Carroll et al., 2010; Kato et al., 2013), and a three-protein complex termed the Mis18 complex (Barnhart et al., 2011; French et al., 2017; Moree et al., 2011; Nardi et al., 2016; Pan et al., 2017; Spiller et al., 2017; Tachiwana et al., 2015). The Mis18 complex recognizes centromeric chromatin either directly by binding to CENP-A nucleosomes (French et al., 2017; Hori et al., 2017; Kral, 2015; Sandmann et al., 2017) or indirectly by binding CCAN components (Dambacher et al., 2012; Moree et al., 2011; Shono et al., 2015; Stellfox et al., 2016). Together, CENP-C and the Mis18 complex restrict HJURP localization to the pre-existing centromere.

The process of CENP-A assembly is restricted to G1 in vertebrates in part by Cdk1 activity (Bernad et al., 2011; Jansen et al., 2007; Moree et al., 2011; Silva et al., 2012). When Cdk activity is inhibited, HJURP and the Mis18 complex localize to centromeres prior to G1 resulting in CENP-A assembly during G2 or even S phase (Muller et al., 2014; Silva et al., 2012). Cdk phosphorylation of HJURP inhibits HJURP binding to Mis18β, preventing its localization (Muller et al., 2014; Stankovic et al., 2016; Wang et al., 2014). Cdk phosphorylation of M18BP1 also inhibits M18BP1 binding to Mis18α/Mis18β and prevents M18BP1 centromere localization in human cells (McKinley and Cheeseman, 2014; Pan et al., 2017; Silva et al., 2012; Spiller et al., 2017; Stankovic et al., 2016). In addition, Cdk1 phosphorylation of CENP-A restricts CENP-A assembly by inhibiting CENP-A association with the HJURP chaperone (Fachinetti et al., 2017; Hu et al., 2011; Yu et al., 2015).

While the Mis18 complex plays a clear role in localizing HJURP to centromeres in G1, whether it has additional roles in regulating CENP-A assembly remains unclear. Notably, M18BP1/KNL2 localizes throughout the cell cycle in *Xenopus* (Moree et al., 2011), *C. elegans* (Maddox et al., 2007), and chicken (Hori et al., 2017; Perpelescu et al., 2015), including all stages of mitosis. Metaphase M18BP1 localization has also been reported in human (McKinley and Cheeseman, 2014). The Mis18 complex has been proposed to ‘prime’ centromeric chromatin for CENP-A assembly, perhaps by regulating centromeric histone acetylation (Fujita et al., 2007; Hayashi et al., 2004; Kim et al., 2012; Ohzeki et al., 2012; Ohzeki et al., 2016; Shang et al., 2016). However, directly tethering HJURP to chromatin is sufficient to drive CENP-A assembly even in the absence of the Mis18 complex (Barnhart et al., 2011; French et al., 2017), suggesting that the primary role of the Mis18 complex in CENP-A assembly is HJURP localization in G1.

To understand how M18BP1 controls CENP-A assembly, we identified the requirements for M18BP1 localization to metaphase and interphase centromeres using a cell-free CENP-A assembly system in *Xenopus laevis* egg extracts. We previously demonstrated that M18BP1 localization to metaphase centromeres in *Xenopus* requires CENP-C (Moree et al., 2011). CENP-C binding by the Mis18 complex is conserved in human, *Xenopus*, and mouse (Dambacher et al., 2012; Moree et al., 2011; Stellfox et al., 2016). Here, we show that the interaction between M18BP1 and CENP-C is mediated by the highly conserved SANTA domain in M18BP1. Their interaction requires M18BP1 phosphorylation at threonine 166, a conserved Cdk site, and is therefore restricted to mitosis. Surprisingly, mutations that disrupt the interaction between M18BP1 and CENP-C not only prevent proper metaphase localization, but also interphase localization. We show that defective interphase localization results in part from defective nuclear localization, raising the possibility that proper metaphase localization promotes retention of M18BP1 in the nucleus during exit from mitosis. We propose that M18BP1 may identify centromeric sites in metaphase for subsequent CENP-A nucleosome assembly in interphase.

## Results

### Metaphase localization of M18BP1 requires the conserved SANTA domain

*Xenopus laevis* expresses two M18BP1 isoforms that are both capable of supporting interphase CENP-A assembly but differ in the timing of their localization to centromeres (Moree et al., 2011; Session et al., 2016). We confirmed that Myc-tagged versions of M18BP1-1 and M18BP1-2 made by *in vitro* translation and added to *Xenopus* egg extract recapitulated this localization to sperm chromatin. Consistent with previous results we found that M18BP1-1 localized to metaphase centromeres while M18BP1-2 was only weakly detectable (Moree et al., 2011) (Supplemental Figure 1A).

To determine which regions of the M18BP1-1 protein were important for metaphase localization we measured the extent of localization of Myc-tagged M18BP1-1 truncations at sperm centromeres. In metaphase extracts depleted of both M18BP1 isoforms, a truncated version of M18BP1-1 containing only the first 580 amino acids (M18BP1-1 ^1-580^) was sufficient for centromeric localization. This result demonstrates that residues 581-1356 of M18BP1-1 - which contain both the CENP-A nucleosome binding domain (French et al., 2017) and the DNA-binding SANT/Myb domain (Maddox et al., 2007) - are dispensable for metaphase localization (Figure 1A–C). A fragment of M18BP1-1 containing amino acids 161-580 retained 67 ± 4% of the protein at centromeres compared to the wild type protein (Figure 1A–C); however, further truncation of this region abolished metaphase localization (Figure 1A–C). Thus, M18BP1-1^161-580^ is sufficient to localize to metaphase centromeres.

**Figure 1:**
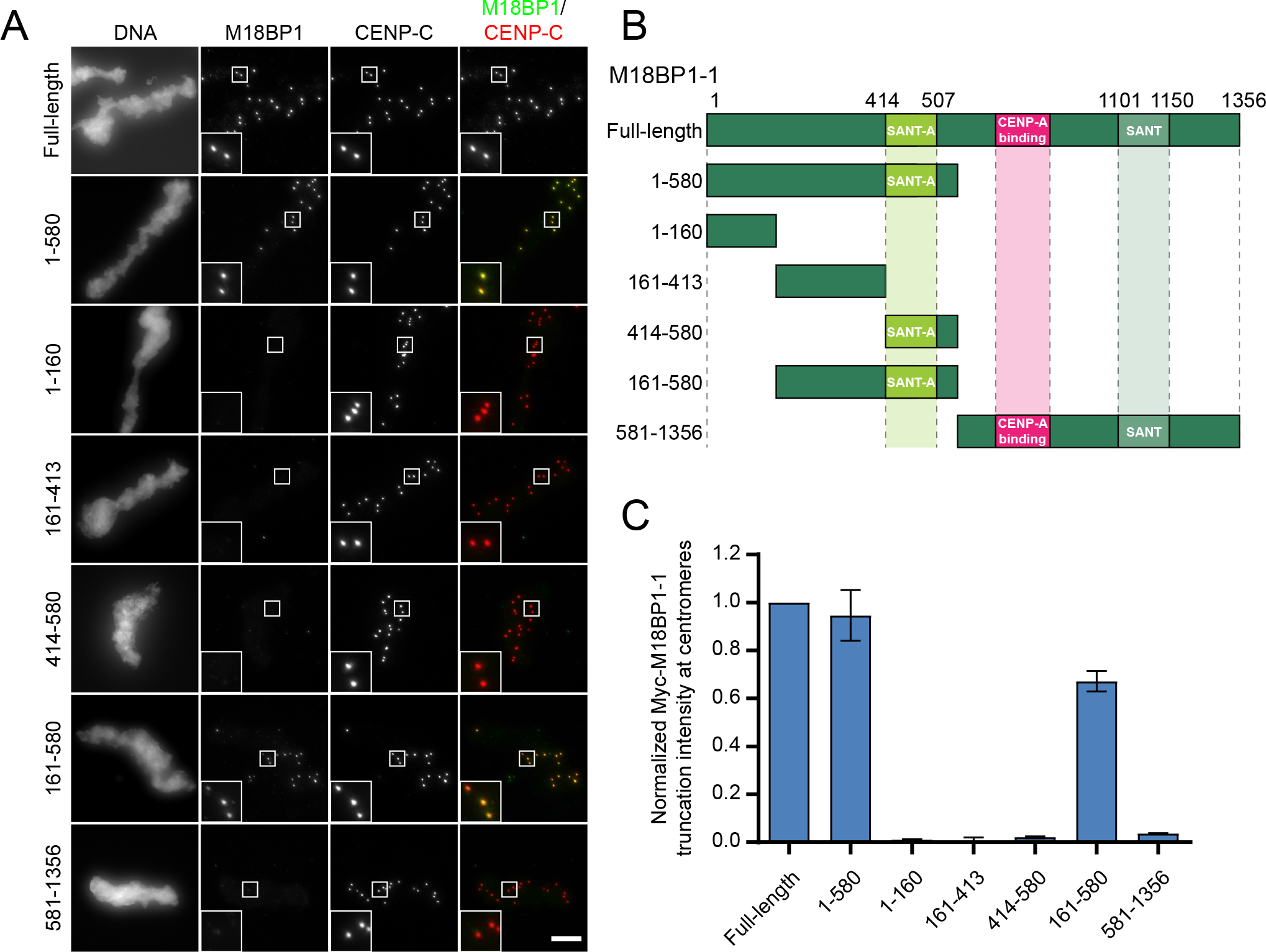
M18BP1-1^161-580^ is sufficient for centromere localization in metaphase egg extract. A) Representative immunofluorescence images showing localization of Myc-tagged M18BP1-1 truncations schematized in (B) to metaphase sperm centromeres in M18BP1-depleted extract. Truncations analyzed are indicated on the left, immunolocalized proteins are indicated above. Scale bar, 10 jm. Insets are magnified 3X. B) Schematic of M18BP1-1 truncations used to define the metaphase targeting domain in (A) and (C). C) Quantification of the average Myc intensity at centromeres from (A). Values are normalized to centromeric signal of full-length M18BP1-1. Graph shows the mean ± SEM of at least three experiments.

M18BP1-1^161-580^ contains a highly conserved SANTA (<u>SANT</u>-<u>A</u>ssociated) domain (Figure 1B), named for its co-occurrence with DNA-binding SANT/Myb domains. The SANTA domain contains five highly conserved hydrophobic residues (Figure 2A) (Zhang et al., 2006). We previously showed that mutation of these residues to alanine in the SANTA domain of M18BP1-2 reduced its localization to interphase centromeres by >90% (French et al., 2017). We tested whether mutation of the SANTA domain in full-length M18BP1-1 also affected metaphase localization and found that M18BP1-1^L416A, W419A, F491A, F495A, W499A^ (M18BP1-1^SANTA^) showed a 45 ± 5% reduction in metaphase localization (Figure 2C–D) indicating that the SANTA domain is required for proper metaphase M18BP1 localization.

**Figure 2:**
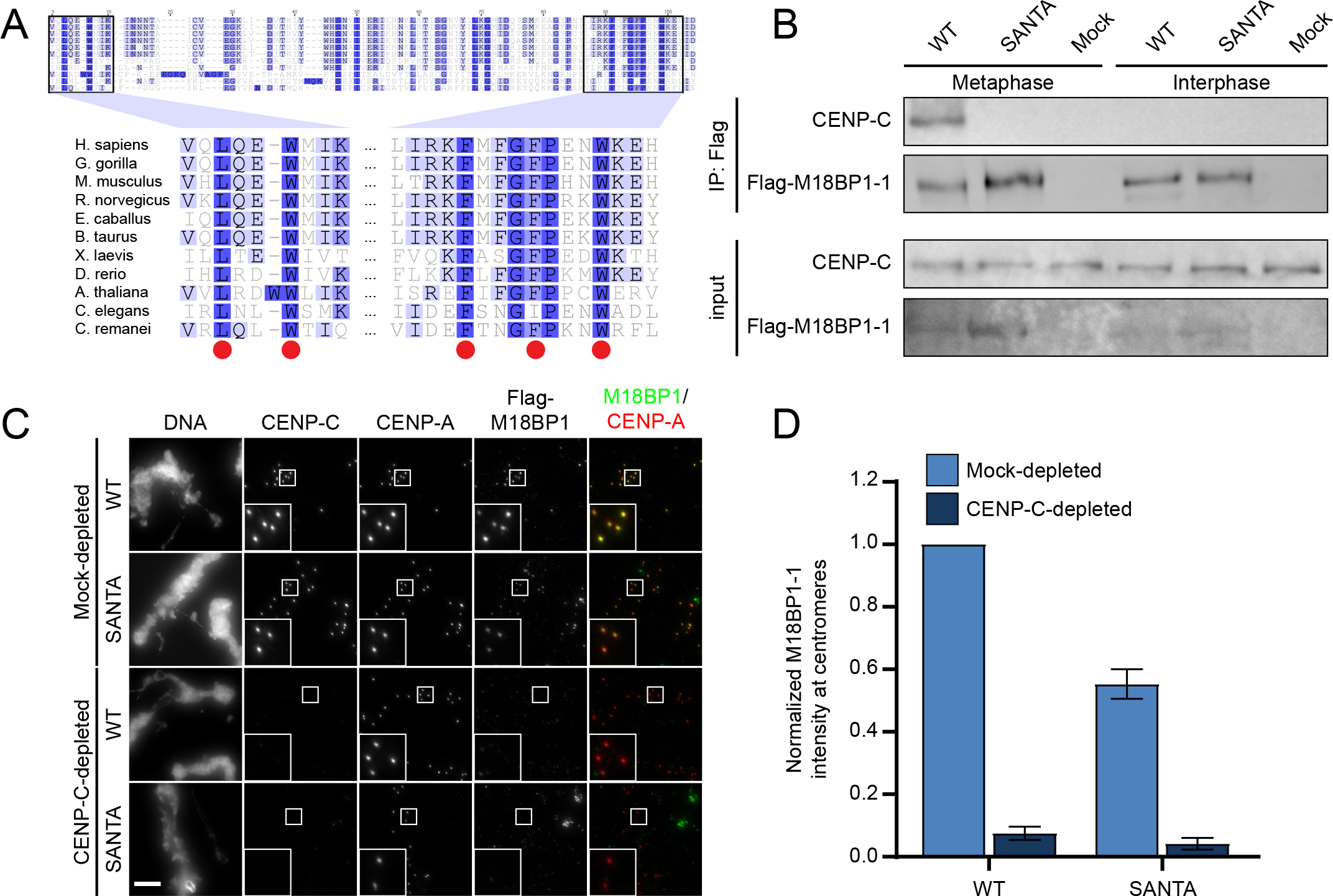
Mutation of the conserved SANTA domain disrupts M18BP1-1 binding to CENP-C and metaphase centromere localization. A) Alignment of SANTA domain among select eukaryotes. Red circles indicate conserved hydrophobic residues predicted to play a role in protein-protein interactions that were mutated in this study (Zhang, et al. 2006). Darker shades of blue represent increased conservation of amino acids in alignment of ~300 M18BP1 homologues. B) Representative Western blot showing disruption of the interaction between M18BP1-1 and CENP-C by mutation of the SANTA domain by co-immunoprecipitation. Extract depleted of endogenous M18BP1 was supplemented with full-length Flag-M18BP1-1^WT^ or Flag-M18BP1-1^SANTA^. The M18BP1-1 species added to each reaction and the cell cycle state of the extract is indicated above. The immunoblotted species is indicated at left. Mock precipitations using rabbit reticulocyte lysate without any *in vitro* translated protein served as a negative control. C) Representative images showing that mutation of the SANTA domain in M18BP1-1 causes decreased localization to sperm centromeres. Metaphase extracts were depleted of endogenous M18BP1 and either mock-depleted or depleted of CENP-C. The extracts were then complemented with WT or SANTA mutant M18BP1-1. M18BP1-1 species and depletion conditions are indicated at left, immunolocalized proteins are indicated above. Scale bar, 10 μm. Insets are magnified 3X. D) Quantification of the average Flag-M18BP1-1 intensity at centromeres from (C). Values are normalized to centromeric signal of M18BP1-1^WT^ in mock-depleted extract. Graph shows the mean ± SEM of three experiments.

*Xenopus* M18BP1 directly interacts with CENP-C, and immunodepletion of CENP-C from egg extract prevents metaphase M18BP1 localization (Moree et al., 2011). We used co-immunoprecipitation to test whether the reduction in centromere localization of the M18BP1-1^SANTA^ mutant was due to a disruption of the interaction between M18BP1-1 and CENP-C. We added *in vitro* translated M18BP1-1^WT^ or M18BP1-1^SANTA^ to metaphase *Xenopus* extracts depleted of endogenous M18BP1. While M18BP1-1^WT^ coprecipitated CENP-C from metaphase extract, M18BP1-1^SANTA^ failed to precipitate CENP-C (Figure 2B) suggesting that the SANTA mutant disrupts CENP-C binding.

We predicted that if metaphase M18BP1 localization were solely mediated by CENP-C binding, then M18BP1-1^SANTA^ localization would not be further reduced following CENP-C depletion. To test the role of CENP-C in localizing M18BP1 we depleted CENP-C from *Xenopus* egg extracts and measured the localization of M18BP1-1^WT^ and M18BP1-1^SANTA^ at centromeres. CENP-C depletion reduced both M18BP1-1^WT^ and M18BP1-1^SANTA^ localization to 8 ± 2% and 4 ± 2% of mock-depleted levels, respectively (Figure 2C–D). Given that the M18BP1-1^SANTA^ mutation prevented co-immunoprecipitation of CENP-C (Figure 2B), these results indicate that either there exists an additional CENP-C-dependent factor required for M18BP1 localization or that the SANTA mutation only partially disrupts the interaction of M18BP1 with CENP-C.

### Amino acids 161-580 comprise the CENP-C binding domain of M18BP1-1

To better characterize the regions of M18BP1 required for CENP-C binding in metaphase, we purified GST-CENP-C^1191-1400^, the previously identified M18BP1-binding domain (Moree et al., 2011), from *E. coli* and performed GST pull-downs with a series of M18BP1-1 truncations expressed by *in vitro* translation (Figure 3A–C, Supplementary Figure 2A). M18BP1-1^1-413^ and M18BP1-1^414-749^, two adjacent but non-overlapping truncations, bound to CENP-C, suggesting that portions of each contribute to the interaction between M18BP1 and CENP-C (Figure 3A–C). M18BP1-1^161-580^, which spans regions of M18BP1-1^1-413^ and M18BP1-1^414-749^, was sufficient to bind CENP-C *in vitro* (Figure 3A–C). We found that the homologous regions in *Xenopus* M18BP1-2 (M18BP1-2^161-570^) and in human M18BP1 (HsM18BP1^325-480^) were also sufficient for CENP-C binding *in vitro* (Supplementary Figures 1B,2B-C). In contrast with a previous report in mouse (Dambacher et al., 2012), we found that the SANT/Myb domain was not required for CENP-C binding (Figure 3A–C).

**Figure 3:**
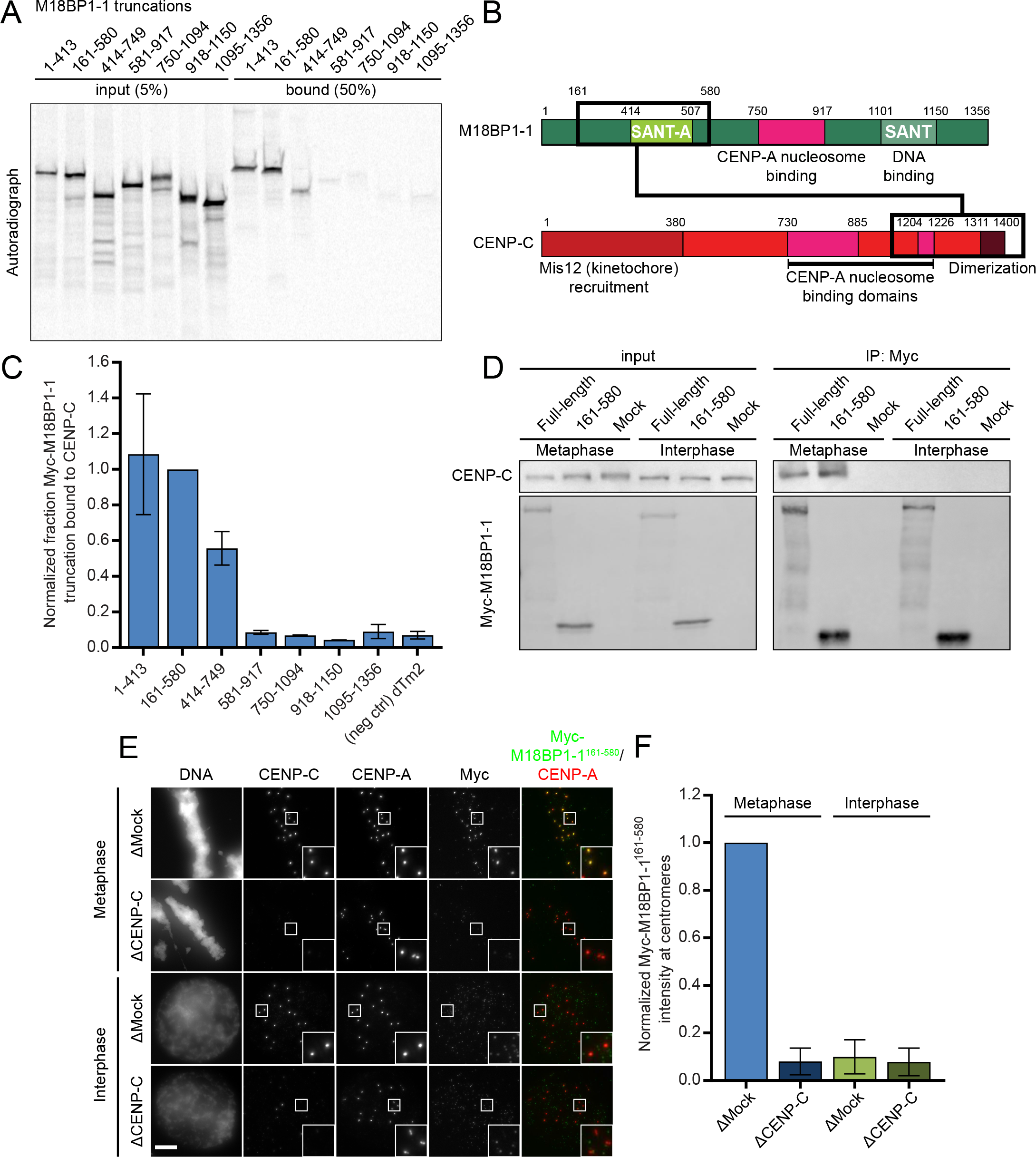
M18BP1-1^161-580^ recapitulates metaphase-specific binding to CENP-C. A) A representative autoradiograph from a GST-pulldown of CENP-C to map the CENP-C binding domain of M18BP1-1. Radiolabeled Myc-M18BP1-1 truncations (amino acids indicated at the top) were mixed with recombinant GST-CENP-C^1191-1400^. Material bound to glutathione agarose was resolved by SDS-PAGE and visualized by autoradiography (see also Supplementary Figure 2A). B) Schematic showing the cognate binding domains on M18BP1 (top) and CENP-C (bottom), indicated by the black boxes, relative to other functional domains. C) Quantification of (A). Bound material as a fraction of the input was calculated from autoradiographs. The graph shows mean fraction bound ± SD of three independent experiments normalized to M18BP1-1^161-580^, the CENP-C binding domain. D) Representative Western blot showing metaphase-specific binding of M18BP1-1^161-580^ to CENP-C by co-immunoprecipitation. Extract depleted of endogenous M18BP1 was supplemented with full-length Myc-M18BP1-1 or Myc-M18BP1-1^161-580^ M18BP1-1 species added to each reaction and cell cycle state of the extract is indicated above. Immunoblotted species indicated at left. Mock precipitations using rabbit reticulocyte lysate without any *in vitro* translated protein served as a negative control. E) Representative immunofluorescence images showing Myc-M18BP1-1^161-580^ localization at sperm centromeres in extract depleted of endogenous M18BP1. In addition, extract was mock-depleted or immunodepleted of CENP-C (indicated at left). Cell cycle state indicated at left, immunolocalized protein indicated above. Scale bar, 10 μm. Insets magnified 3X. F) Quantification of (E). Graph shows mean Myc-M18BP1-1^161-580^ intensity at centromeres ± SEM of two independent experiments normalized to the mock-depleted, metaphase condition.

M18BP1-1 binds directly to CENP-C in metaphase, but not in interphase (Moree et al., 2011). To test whether M18BP1-1^161-580^ retained the cell cycle dependent interaction with CENP-C in *Xenopus* egg extract, we added M18BP1-1^161-580^ to metaphase and interphase extract, immunoprecipitated the protein and assayed CENP-C coprecipitation by Western blotting. Similar to full-length M18BP1-1, we found that M18BP1-1^161-580^ co-precipitates CENP-C only from metaphase egg extract (Figure 3D). We next assayed the localization of M18BP1-1^161-580^ to sperm centromeres in egg extract and found that M18BP1-1^161-580^ associates only with metaphase centromeres. Immunodepletion of CENP-C from these extracts prevented the localization of M18BP1-1^160-580^ to the centromere (Figure 3E–F). Thus, M18BP1-1^161-580^ binds directly to CENP-C, co-precipitates CENP-C in metaphase, and localizes to metaphase centromeres dependent on the presence of CENP-C.

### CDK phosphorylation of threonine 166 is required for metaphase M18BP1-1 localization and CENP-C binding

During the transition from metaphase to interphase, M18BP1 switches from binding directly to CENP-C for metaphase centromere localization to binding directly to CENP-A nucleosomes for interphase centromere localization (French et al., 2017; Moree et al., 2011). We tested whether the interaction between M18BP1 and CENP-C might be regulated by a shift in mitotic kinase activity as cells exit metaphase. We immunoprecipitated either full-length M18BP1-1 or M18BP1-1^161-580^ from metaphase extract and treated the immunoprecipitate with λ-phosphatase. Both M18BP1 and CENP-C migrated more rapidly in SDS-PAGE gels after λ-phosphatase treatment, consistent with dephosphorylation (Figure 4A–B). Dephosphorylation by λ-phosphatase caused dissociation of CENP-C from both full-length M18BP1-1 and M18BP1-1^161-580^ precipitates (Figure 4A–B). This suggests that phosphorylation is required to maintain the interaction between M18BP1 and CENP-C, and that regulation of this interaction is preserved in M18BP1-1^161-580^.

**Figure 4:**
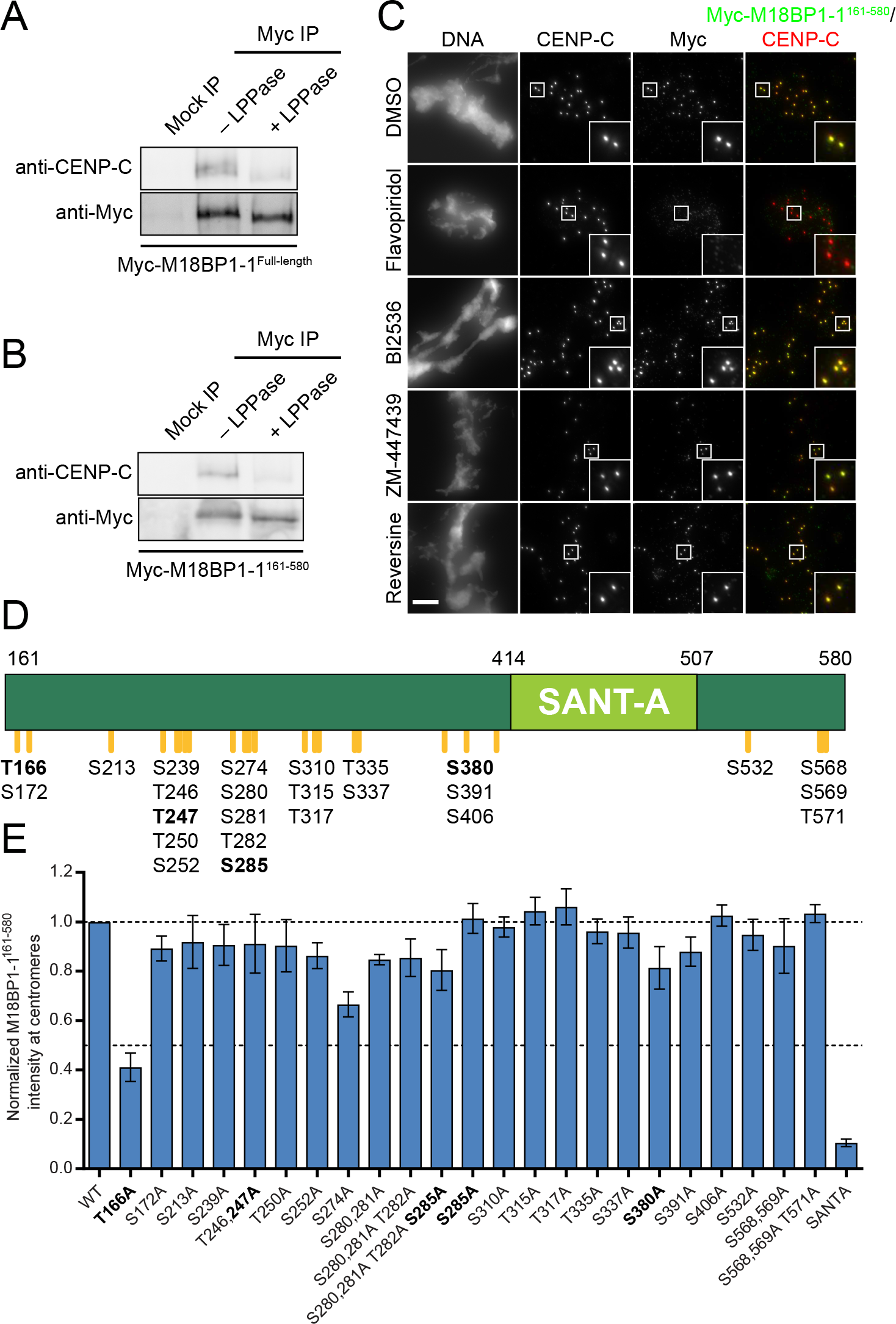
Identification of metaphase phosphorylation sites in M18BP1-1 by mass spectrometry. A-B) Metaphase binding of M18BP1-1 to CENP-C requires phosphorylation. Either full-length Myc-M18BP1-1 (A) or Myc-M18BP1-1^161-580^ (B) was immunoprecipitated from metaphase extract. Half of the immunoprecipitate was treated with A-protein phosphatase (+ LPPase) to assess whether binding to CENP-C required phosphorylation. Immunoprecipitation with an equivalent amount of mouse IgG served as a negative control. Immunoblotted species is indicated at left. C) Cdk inhibition with flavopiridol prevents M18BP1-1^161-580^ localization at metaphase centromeres. Representative images showing Myc-M18BP1-1^161-580^ localization at sperm centromeres in metaphase extract depleted of endogenous M18BP1 following treatment with various mitotic kinase inhibitors. Inhibitor treatment is indicated at left, immunolocalized protein is indicated above. Scale bar, 10 μm. Insets are magnified 3X. D) Schematic of phosphorylation sites in M18BP1-1^161-580^ mutated for this study. M18BP1-1 residue numbers indicated above. Positions of phosphorylation sites are indicated by orange lines and labels below. Bold indicates consensus Cdk phosphorylation sites. E) Quantification of immunofluorescence experiments examining localization of M18BP1-1^161-580^ phosphorylation site mutants to sperm centromeres in metaphase extract depleted of endogenous M18BP1. Graph shows mean centromere intensity ± SEM of three independent experiments normalized to the WT condition. See also Supplemental Figure 4B.

To identify kinases that might regulate M18BP1 localization, we analyzed M18BP1-1^161-580^ localization in M18BP1-depleted metaphase extract treated with inhibitors of several mitotic kinases: cyclin-dependent kinase (Cdk; flavopiridol), Polo-like kinase (Plx; BI2536), Aurora kinase (ZM-447439), and Mps1 (reversine). Of these, only Cdk inhibition caused loss of M18BP1-1^161-580^ localization (Figure 4C). Although, Cdk inhibition also drives the extract into interphase, inhibitors of other mitotic kinases had no effect on M18BP1 localization suggesting that Cdk phosphorylation may directly regulate M18BP1.

Because M18BP1-1^161-580^ is phosphorylated in metaphase extract (Figure 4B) we tested whether M18BP1 phosphorylation might regulate its centromere localization by modulating its interaction with CENP-C. We purified MBP-tagged M18BP1-1^161-580^ from *E. coli* and added the purified protein to M18BP1-depleted egg extract (Supplemental Figure 3). We then recovered the protein from the extract with α-MBP antibody-coated beads and mapped modification sites on M18BP1-1^161-580^ with mass spectrometry (Figure 4D, Supplemental Figure 3). We identified 25 phosphorylated residues, including four minimal consensus Cdk sites (S/T-P) (Figure 4D). We mutated each of these phosphorylation sites to alanine to prevent their modification and assessed the localization of mutant M18BP1-1^161-580^ at sperm centromeres in metaphase extract depleted of endogenous M18BP1. Most mutants exhibited a modest (~10%) reduction in metaphase localization (Figure 4E). However, mutation of one conserved Cdk site, T166A, caused a 59 ± 6% decrease in M18BP1-1^161-580^ localization (Figure 4E, Supplemental Figure 4B). Mutation of the SANTA domain caused an 89 ± 2% decrease in this assay (Figure 4E).

When introduced into the full length M18BP1 protein, M18BP1-1^T166A^ caused a substantial 77 ± 3% reduction in metaphase localization whereas two control mutants, M18BP1-1^S172A^ and M18BP1-1^S568A, S569A^, showed only modest localization defects (Figure 5A,C, Supplemental Figure 4C–D). This indicates that Cdk phosphorylation of T166 is primarily responsible for regulating metaphase M18BP1 localization. To test if the centromere localization defect in M18BP1-1^T166A^ was due to loss of interaction with CENP-C, we immunoprecipitated M18BP1-1 mutants from metaphase extract and found that M18BP1-1^WT^, M18BP1-1^S172A^, and M18BP1-1^S568A, S569A^ all co-precipitated CENP-C while M18BP1-1^T166A^ did not (Figure 5B). This indicates that the defect in M18BP1-1^T166A^ localization to metaphase centromeres likely arises from the disruption of its interaction with CENP-C.

**Figure 5:**
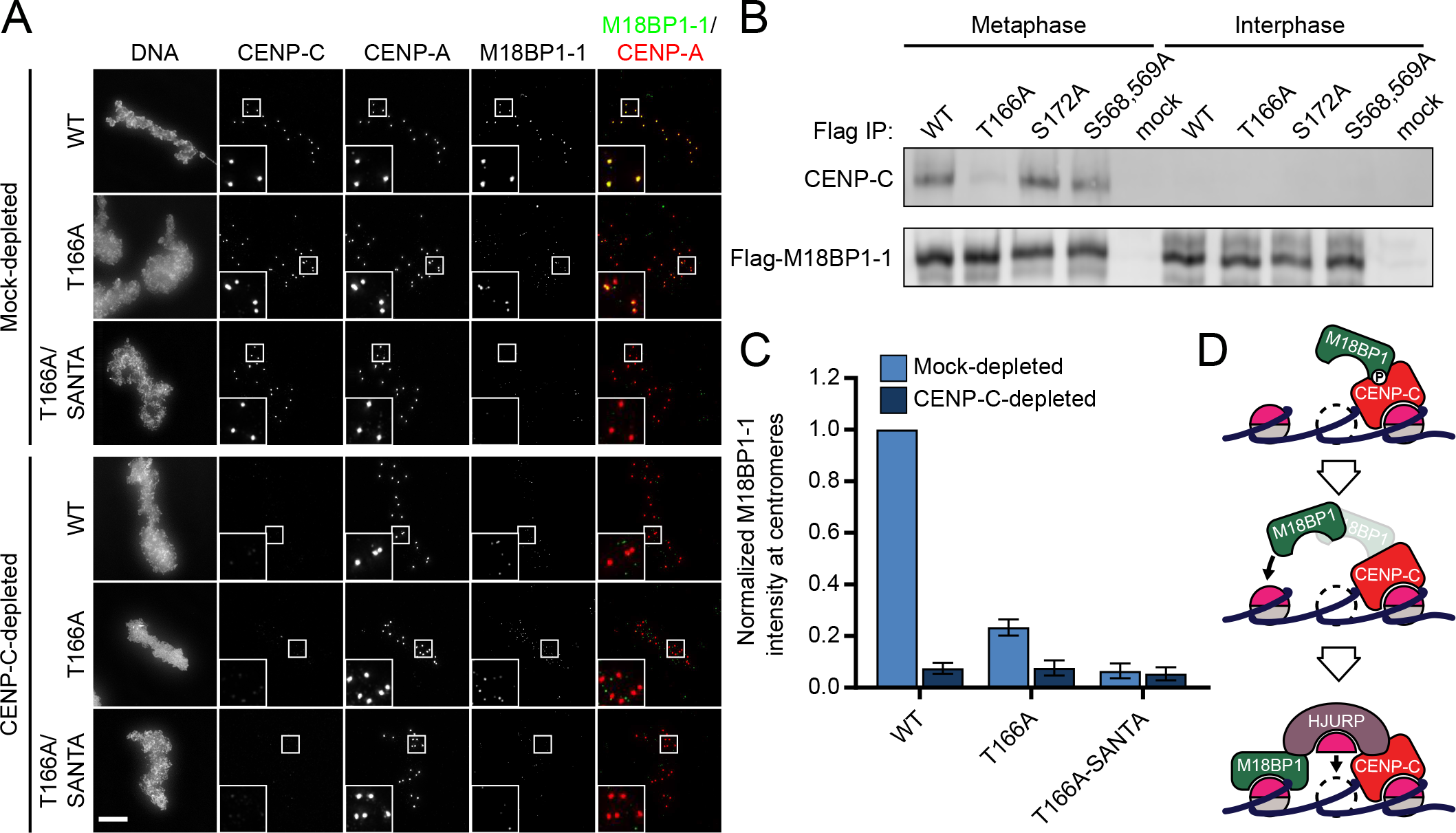
Phosphorylation of T166 in M18BP1 regulates metaphase localization and binding to CENP-C. A) Representative immunofluorescence images showing M18BP1-1 mutant localization. M18BP1-depleted metaphase extract was supplemented with *in vitro* translated full-length Flag-M18BP1-1 mutants and sperm chromatin. In addition, extract was either mock-depleted or CENP-C depleted to assess CENP-C dependent metaphase localization. M18BP1-1 species and CENP-C depletion status are indicated at left, immunolocalized protein is indicated above. Scale bar, 10 μm. Insets are magnified 3X. B) Representative Western blot showing co-immunoprecipitation of CENP-C with phosphorylation site mutants of M18BP1-1. M18BP1-depleted metaphase or interphase extract was supplemented with *in vitro* translated Flag-M18BP1-1 mutants. M18BP1-1 mutation and cell cycle state indicated above, immunoblotted species indicated at right. Mock precipitations using rabbit reticulocyte lysate without any *in vitro* translated protein served as a negative control. C) Quantification of (A). Flag-M18BP1-1 mutant centromere intensity normalized to WT localization in the mock-depleted condition. Error bars represent SEM from three independent experiments. D) Model showing phosphorylation-dependent binding of M18BP1 to CENP-C during mitosis (top) and the transition to CENP-A-dependent binding in interphase (bottom).

We determined the extent to which M18BP1-1 depends on its interaction with CENP-C for localization to metaphase centromeres by depleting CENP-C from egg extract and assaying the localization of the M18BP1-1^T166A^ mutant. M18BP1-1^T166A^ localization was further reduced by CENP-C depletion to 8 ± 3% of mock-depleted levels (Figure 5A, C). This, again, indicated that either M18BP1-1^T166A^ mutation does not completely disrupt the interaction between M18BP1 and CENP-C or that there exist additional, CENP-C-dependent factor(s) required for M18BP1 localization.

Our prior *in vitro* binding experiments suggested that separate elements contained within M18BP1-1^1-413^ or M18BP1-1^414-749^ were sufficient to bind to CENP-C (Figure 3A–B). Given that M18BP1-1^1-413^ contains T166 but lacks the SANTA domain, and that M18BP1-1^414-749^ contains the SANTA domain but lacks T166, we reasoned that the T166A or SANTA mutations in isolation might not completely ablate CENP-C binding. Combining these mutations in a M18BP1-1^T166A, SANTA^ double mutant completely abolished metaphase M18BP1-1 localization, comparable to that observed after CENP-C depletion (Figure 5A, C). This indicates not only that both contacts contribute to CENP-C binding, but also suggests that metaphase M18BP1 localization is mediated exclusively by binding to CENP-C (Figure 5D).

### M18BP1-1^T166A^ and M18BP1-1^SANTA^ do not support proper CENP-A assembly due to defective interphase localization

CENP-C depletion prevents proper CENP-A assembly in interphase in part because CENP-C directly recruits HJURP to centromeres (French et al., 2017; Moree et al., 2011; Tachiwana et al., 2015). Although CENP-C is not required for interphase M18BP1 localization, it was unclear whether mislocalization of M18BP1 in metaphase might also alter the efficiency of CENP-A assembly. To uncover a possible requirement for metaphase M18BP1 localization, we asked whether M18BP1 mutants defective in metaphase localization could support interphase CENP-A assembly.

To assess this, we depleted endogenous M18BP1 from interphase extract, replaced it with *in vitro* translated M18BP1-1 mutants, and assayed the assembly of exogenously added Myc-CENP-A at sperm centromeres (Figure 6A–C). Consistent with previous results, we found that M18BP1 depletion reduced CENP-A assembly to 39 ± 3% of mock-depleted levels (Figure 6A–B). Add-back of both M18BP1 isoforms is necessary to fully rescue CENP-A assembly on sperm chromatin (French et al., 2017; Moree et al., 2011). However, addition of only wild-type M18BP1-1 partially rescued assembly to 62 ± 3% of mock-depleted levels (Figure 6A–B). Add-back of M18BP1-1^T166A^ failed to rescue Myc-CENP-A assembly to the same degree, achieving only 53 ± 4% of mock-depleted levels, and add-back of M18BP1-1^SANTA^ rescued Myc-CENP-A assembly to 44 ± 3% of mock-depleted levels (French et al., 2017) (Figure 6A–B). Thus, M18BP1 mutants defective in metaphase localization do not fully rescue interphase CENP-A assembly.

**Figure 6:**
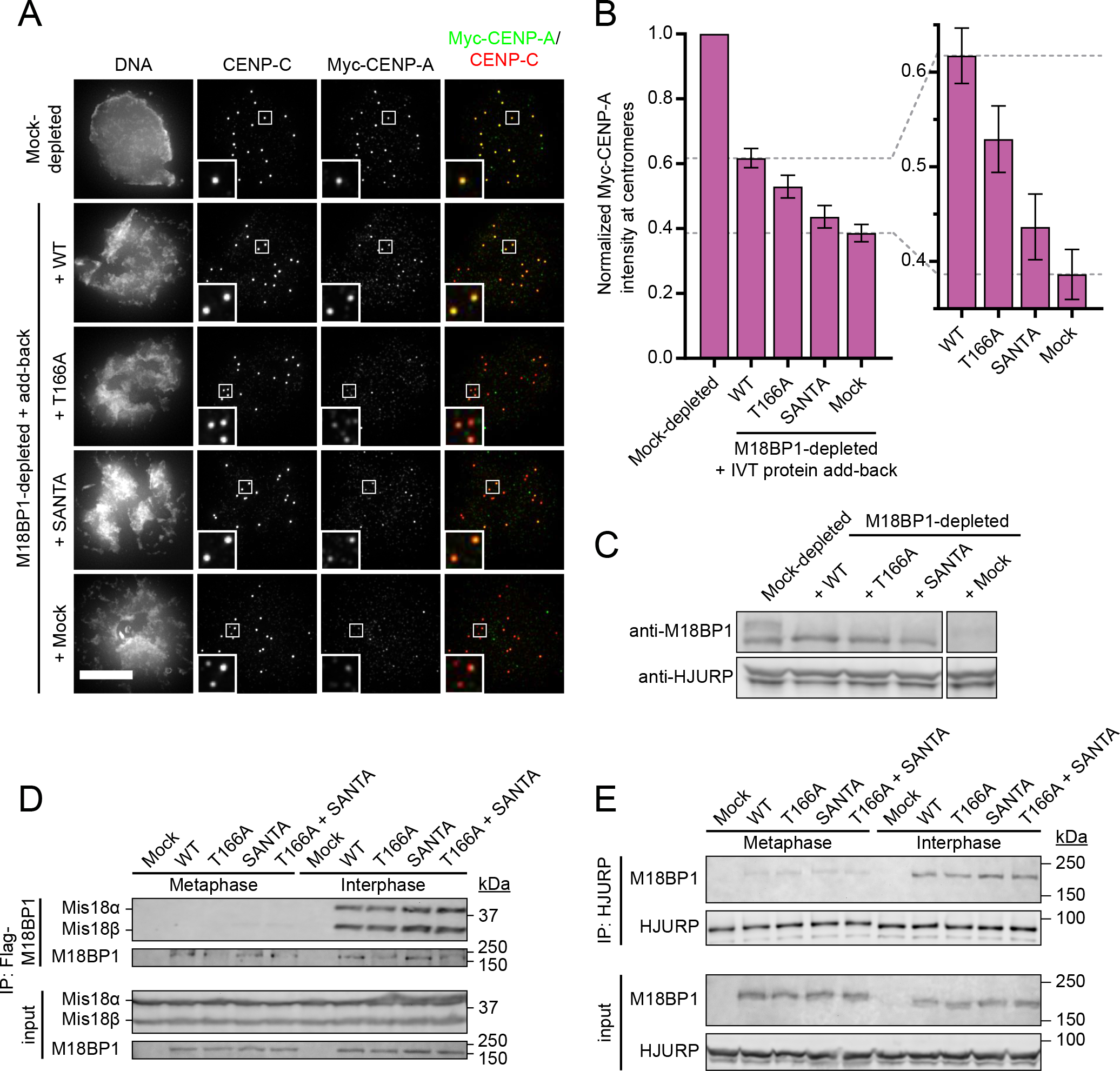
Metaphase targeting mutants of M18BP1-1 do not support robust CENP-A assembly. A) Representative immunofluorescence images showing incorporation of exogenous Myc-CENP-A at centromeres. Sperm nuclei were incubated in M18BP1-depleted interphase *Xenopus* egg extracts complemented with the indicated M18BP1-1 protein. Extracts were supplemented with RNA encoding Myc-CENP-A to track new CENP-A assembly and *in vitro* translated HJURP. Scale bar, 10 μm. Insets are magnified 3X. B) Quantification of Myc-CENP-A loading at Xenopus sperm centromeres in (A). Values are normalized to the centromere signals in mock-depleted extract. Dashed lines indicate the Myc-CENP-A assembly signal observed upon M18BP1 depletion (bottom) and Flag-M18BP1-1^WT^ add-back (top) as points of reference for mutant rescue. Graph shows the mean ± SEM of four independent experiments. C) Representative Western blot of CENP-A assembly reactions in (A) probed with α-M18BP1 (top) and α-HJURP (bottom). Efficient M18BP1 depletion is indicated by comparing lanes 1 and 5. Add-back of wild-type or mutant M18BP1-1 is near endogenous levels. D) Interphase Mis18 complex formation is unaffected by M18BP1-1 mutations that prevent binding to CENP-C. Extract depleted of endogenous M18BP1 was supplemented with Myc-Mis18α, Myc-Mis18β, and Flag-M18BP1-1. M18BP1-1 species added to each reaction and cell cycle state of the extract is indicated at the top. Co-immunoprecipitation of Myc-Mis18α/β was assessed by α-Myc immunoblot following Flag precipitation. E) HJURP association with M18BP1-1 is unaffected by mutations that prevent CENP-C binding. Extract depleted of endogenous M18BP1 was supplemented with Flag-M18BP1-1. M18BP1-1 species added to each reaction and cell cycle state of the extract are indicated at the top. Coimmunoprecipitation of Flag-M18BP1-1 was assessed by α-Flag immunoblot following HJURP precipitation.

M18BP1, as part of the Mis18 complex, recruits HJURP to centromeric chromatin for CENP-A nucleosome assembly (Barnhart et al., 2011; French et al., 2017; Moree et al., 2011; Nardi et al., 2016). To understand why metaphase targeting mutants of M18BP1 were defective in Myc-CENP-A assembly, we examined their association with Mis18α, Mis18β, and HJURP by immunoprecipitation followed by Western blotting. We observed interphase-specific association between *Xenopus* Mis18α, Mis18β, and M18BP1-1^WT^ by immunoprecipitation (Figure 6D). M18BP1-1^T166A^, M18BP1-1^SANTA^ and M18BP1-1^T166A,SANTA^ bound to Mis18α and Mis18β similar to wild-type, suggesting that defects in CENP-A assembly do not result from defective Mis18 complex assembly. We also assayed binding to HJURP and found that M18BP1-1^WT^, M18BP1-1^T166A^, M18BP1-1^SANTA^ and M18BP1-1^T166A,SANTA^ all bound to HJURP in interphase (Figure 6E). These data suggest that M18BP1-1 mutants that are defective in metaphase centromere localization are not compromised in their ability to assemble the Mis18 complex or interact with HJURP.

We had previously found that mutation of the SANTA domain in M18BP1-2 prevented its localization in interphase (French et al., 2017). Similarly, we found that M18BP1-1^SANTA^ localization to interphase centromeres was reduced by 79 ± 5% (Figure 7A–B). Surprisingly, we also found that interphase localization of M18BP1-1^T166A^ was reduced by 54 ± 6% (Figure 7A–B). Defective interphase localization of these mutants is the likely cause of reduced Myc-CENP-A assembly because M18BP1 localization to centromeres in interphase is required for new CENP-A nucleosome assembly.

**Figure 7:**
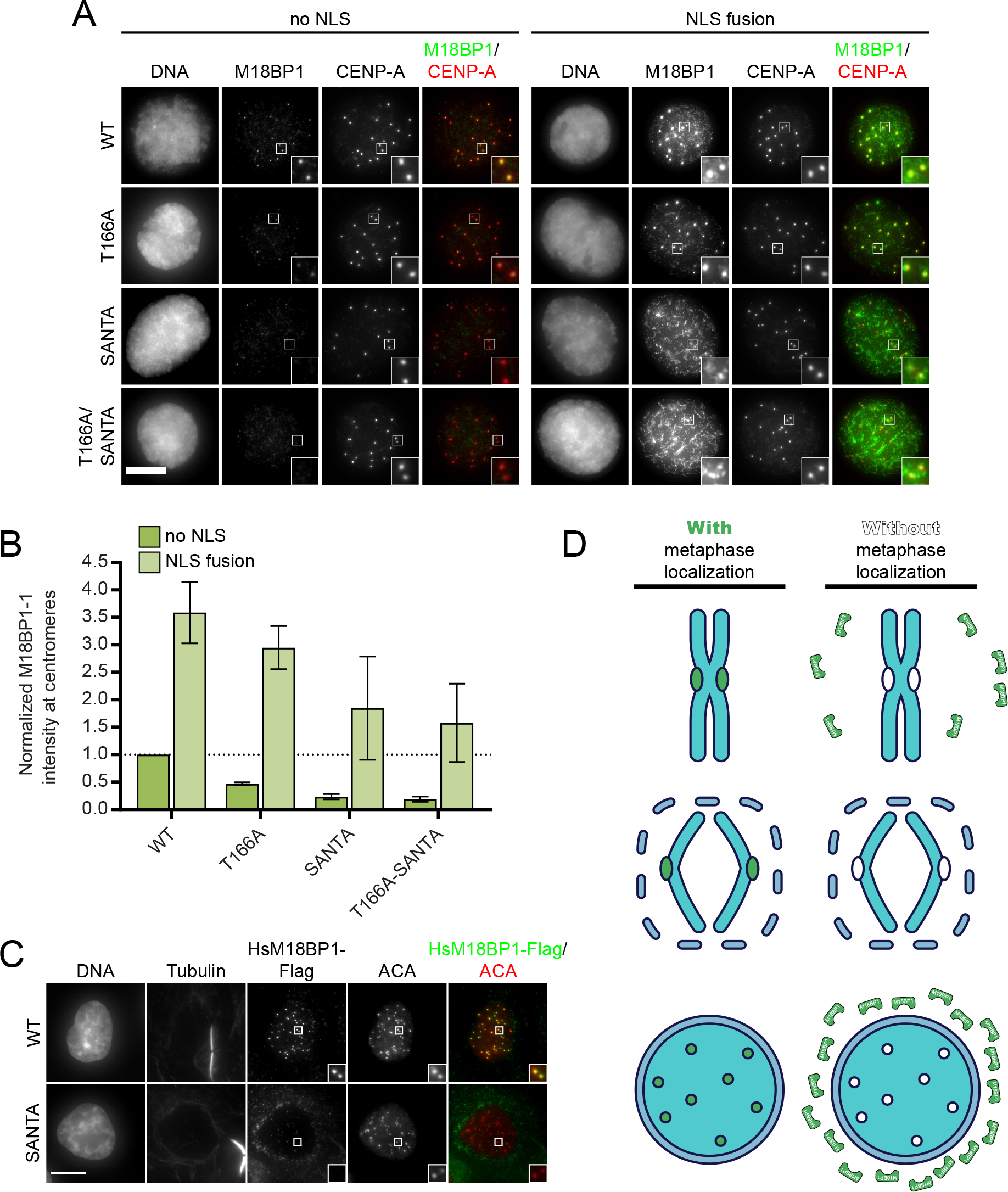
Metaphase targeting mutants of M18BP1-1 inhibit nuclear localization during interphase. A) Representative immunofluorescence images showing localization of full-length Flag-M18BP1-1 (mutant species indicated at left) without (left panels) and with (right panels) fusion to an SV40 NLS to sperm centromeres in interphase extract depleted of endogenous M18BP1. Immunolocalized protein indicated above. Scale bar, 10 μm. Insets are magnified 3X. B) Quantification of immunofluorescence intensity of full-length Flag-M18BP1-1 (mutant species indicated at bottom) at interphase centromeres from (A). Graph shows mean centromere intensity normalized to WT with no NLS. Error bars represent SEM from three independent experiments. C) Representative images showing G1 localization of human M18BP1^WT^ or human M18BP1^SANTA^ in sfGFP-AID-M18BP1 DLD1 cells treated with 1 mM IAA for 24 h to remove endogenous M18BP1. Centromeric localization is indicated by localization with ACA (α-centromere autoantibody serum), early G1 cell cycle state is indicated by midbody staining in the tubulin channel. M18BP1 species indicated at left, immunolocalized protein indicated above. Scale bar, 10 um. Insets are magnified 3X. D) Model showing that metaphase localization of M18BP1 (green) promotes M18BP1 retention on chromosomes during nuclear envelope formation to promote centromeric localization during interphase.

Prior work has identified only two mechanisms responsible for interphase M18BP1 localization: binding to Mis18α/β and binding to CENP-A nucleosomes (French et al., 2017; Fujita et al., 2007; Hori et al., 2017; Pan et al., 2017; Sandmann et al., 2017; Spiller et al., 2017). As described above, immunoprecipitation of Flag-M18BP1-1 mutants from extract indicated that interphase-specific binding to Mis18α/β was intact (Figure 6D). Mutation of T166A or the SANTA domain did not prevent M18BP1 binding to CENP-A chromatin-coated beads *in vitro*, confirming that CENP-A nucleosome binding was not dramatically impaired (Supplementary Figure 5) (French et al., 2017). This suggests that an additional mechanism may act through the CENP-C binding domain to recruit M18BP1 to interphase centromeres.

### The SANTA domain plays a conserved role in interphase M18BP1 localization in humans

In contrast to our findings in *Xenopus*, deletion of the SANTA domain from human or *Arabidopsis* M18BP1 did not alter M18BP1 localization in these organisms (Lermontova et al., 2013; Stellfox et al., 2016). To understand the apparent difference in SANTA domain function between human cells and *Xenopus*, we generated human cell lines expressing the F-box protein TIR1 and in which both alleles of M18BP1 were endogenously tagged with a tandem sfGFP-AID (auxin-inducible degron) tag, permitting rapid, inducible degradation for complementation studies (Supplementary Figure 6A–C) (Holland et al., 2012).

Visualization of endogenous sfGFP-AID-M18BP1 signal revealed punctate, centromeric localization in midbody-positive G1 cells, consistent with previous reports (Supplementary Figure 6C) (Fujita et al., 2007; Maddox et al., 2007; McKinley and Cheeseman, 2014). We did not observe substantial M18BP1 localization at centromeres during mitosis until anaphase/telophase (Supplementary Figure 6D). Upon addition of 1 mM indole-3-acetic acid (IAA), M18BP1 protein levels were depleted within 15 minutes, and centromeric M18BP1 was undetectable at G1 centromeres within 1 hour (Supplementary Figure 6B–C).

Ectopic expression of wild-type M18BP1 from a doxycycline-inducible promoter restored M18BP1 signal at G1 centromeres (Figure 7C, Supplementary Figure 6E). Expression of M18BP1^SANTA^, however, showed diffuse cytoplasmic localization and appeared to be excluded from the nucleus (Figure 7C, Supplementary Figure 6E), supporting a conserved role for the SANTA domain in interphase M18BP1 localization in human cells and frogs. Centromeric M18BP1^SANTA^ localization was not observed in any stage of the cell cycle. This demonstrates that the SANTA domain plays a conserved, CENP-C-independent role in ensuring the localization of M18BP1 to interphase centromeres in humans and in *Xenopus.*

### A role for the nuclear envelope in regulating M18BP1 localization

The diffuse, cytoplasmic localization of M18BP1^SANTA^ in human cells suggested a defect in nuclear localization (Figure 7C). Given that the T166A and SANTA mutations in *Xenopus* M18BP1-1 did not prevent binding to CENP-A nucleosomes or to Mis18α/β, we hypothesized that their interphase localization defects resulted from defective nuclear localization.

To test if nuclear localization influences M18BP1 targeting, we fused an SV40 nuclear localization sequence (NLS; PKKKRKV) to M18BP1-1 in order to bypass nuclear localization defects. Flag-NLS-M18BP1-1^WT^ showed 3.6-fold increase in localization at interphase centromeres relative to Flag-M18BP1-1^WT^ lacking an NLS (Figure 7A–B). Strikingly, Flag-NLS-M18BP1-1^T166A^ localization increased 6.3-fold relative to Flag-M18BP1-1^T166A^ lacking an NLS, achieving localization comparable to Flag-NLS-M18BP1^WT^ (Figure 7A–B). Flag-NLS-M18BP1-1^T166A^ localization was punctate and largely resembled Flag-NLS-M18BP1-1^WT^ localization. Together, this suggests that the T166A mutation may not directly interfere with interphase M18BP1-1 recruitment to centromeres, but may indirectly prevent it by inhibiting nuclear localization (Figure 7D).

Flag-NLS-M18BP1-1^SANTA^ and Flag-NLS-M18BP1-1^T166A,SANTA^ showed improved nuclear localization and increased signal at centromeres (Figure 7A–B). Intriguingly, however, their localization was not centromere-specific, as was observed for Flag-NLS-M18BP1-1^WT^ or Flag-NLS-M18BP1-1^T166A^: Flag NLS-M18BP1-1^SANTA^ signal was observed throughout chromatin with enrichment around centromeres (Figure 7A). This suggests that the SANTA domain plays a dual role, promoting both nuclear retention of M18BP1-1 during mitotic exit and forming additional interactions that are required for interphase centromere localization.

## Discussion

Maintaining one centromere per chromosome across generations is important for faithful genome transmission. This process requires the assembly of CENP-A nucleosomes every cell cycle, but how CENP-A assembly factors are recruited to the right locus and how CENP-A assembly is regulated are incompletely understood. An emerging theme has been that Cdk phosphorylation inhibits premature assembly, and that dephosphorylation promotes the localization and function of proteins required for CENP-A assembly. Here, we show that Cdk phosphorylation controls the interaction between M18BP1 and CENP-C to promote metaphase centromere localization of M18BP1 in *Xenopus* egg extract. The interaction between CENP-C and M18BP1 is through the C-terminal region of CENP-C containing the conserved CENP-C motif and the cupin/dimerization domain (Moree et al., 2011), and the N-terminal region of M18BP1 containing the conserved SANTA domain. Mutations in M18BP1 that break this interaction prevented metaphase M18BP1 localization, suggesting that CENP-C binding is the primary mechanism for metaphase M18BP1 localization (Figure 5D).

At anaphase, Cdk activity declines. This causes a switch in the mechanism of M18BP1 localization between metaphase and interphase: M18BP1 no longer requires CENP-C, but rather binds to CENP-A nucleosomes for interphase centromere localization in *Xenopus* (Figure 5D) (French et al., 2017; Moree et al., 2011). While CENP-C depletion does not prevent interphase M18BP1 localization, M18BP1 mutants unable to bind CENP-C showed reduced interphase localization, due in part to defective nuclear localization. This apparent paradox may partly be explained by the fact that CENP-C occupies potential M18BP1 binding sites on CENP-A chromatin (French et al., 2017); the liberation of additional M18BP1 binding sites by CENP-C depletion may allow ‘extra’ M18BP1 to bind centromeres before nuclear envelope assembly. Our data are consistent with a model in which metaphase localization of M18BP1, or at least chromatin association prior to nuclear envelope reformation, promotes proper interphase localization and therefore CENP-A assembly (Figure 7D).

Metaphase localization of M18BP1 could serve additional functions beyond merely ensuring its retention during nuclear assembly. In interphase, both CENP-C and M18BP1 are required for HJURP localization and CENP-A assembly (French et al., 2017; Moree et al., 2011). Metaphase binding of M18BP1 to CENP-C could ensure their close apposition in the subsequent interphase on adjacent nucleosomes or even opposite faces of the same CENP-A nucleosome, suggesting a mechanism by which sites for new CENP-A assembly might be identified prior to G1. Their interaction could also provide an additional layer of regulation to prevent premature HJURP localization. HJURP interacts with CENP-C^1191-1400^ during interphase (French et al., 2017; Tachiwana et al., 2015). Binding of M18BP1 to this region in metaphase could competitively inhibit HJURP localization, helping to restrict CENP-A assembly to G1 (Figure 5D).

While our work demonstrated a role for the SANTA domain in regulating M18BP1 nuclear localization, our inability to rescue centromere-specific localization of M18BP1-1^SANTA^ by fusion to an NLS suggests the domain regulates additional aspects of interphase localization. The importance of the SANTA domain is further highlighted by the fact that preventing CENP-A nucleosome binding by M18BP1 only reduced interphase M18BP1 localization by ~50% (French et al., 2017). Therefore, a major goal moving forward will be to identify the interphase functions of the SANTA domain to understand how the Mis18 complex recognizes CENP-A chromatin during CENP-A assembly.

Our work also suggests that M18BP1 regulates CENP-A assembly similarly in frog and in human. Although previous work found that deletion of the SANTA domain from full-length M18BP1 did not prevent centromere localization in human cells, we identified a conserved role for the SANTA domain in G1 localization of human M18BP1. Two copies of M18BP1 are present in the Mis18 complex (Pan et al., 2017; Spiller et al., 2017), so the presence of endogenous, wild-type M18BP1 may have supported localization of SANTA-deleted M18BP1 (Stellfox et al., 2016). In addition, we found that human M18BP1 interacts directly with CENP-C via a domain homologous to that in *Xenopus*, suggesting conservation of this pathway. Indeed, CENP-C-dependent, metaphase localization of M18BP1 has been previously observed (McKinley and Cheeseman,. Metaphase localization may be transient or highly dynamic in humans, however, as centromeric M18BP1 signal remains low prior to anaphase (Fujita et al., 2007; Lagana et al., 2010; Maddox et al., 2007; Nardi et al., 2016; Silva et al., 2012; Spiller et al., 2017; Stankovic et al., 2016; Wang et al., 2014). The cell free system we have developed in *Xenopus* egg extract provides a powerful approach for investigating the conserved cell cycle dynamics of M18BP1 to understand how the Mis18 complex regulates CENP-A assembly.

## Materials and Methods

### Experimental model and subject details

Mature *Xenopus laevis* females (Nasco, LM00535MX) were housed and maintained in the Stanford Aquatic Facility staffed by the Veterinary Service Center. For ovulation, frogs were primed 2-14 days before ovulation by subcutaneous injection of 50 U pregnant mare serum gonadotropin (PMSG; Sigma) at the dorsal lymph sac, then induced 18 hours before ovulation with 500 U human chorionic gonadotropin (hCG; Chorulon). During ovulation, frogs were housed individually in 2 L MMR buffer (6 mM Na-HEPES, pH 7.8, 0.1 mM EDTA, 100 mM NaCl, 2 mM KCl, 1 mM MgCl_2_, and 2 mM CaCl_2_) at 17°C. Animal work was carried out in accordance with the guidelines of the Stanford University Administrative Panel on Laboratory Animal Care (APLAC).

Human DLD1 cell lines (male, pseudo-diploid, colorectal adenocarcinoma) stably expressing the F-box protein osTIR1 were a gift from Don Cleveland (Ludwig Cancer Institute, San Diego) (Fachinetti et al., 2015; Holland et al., 2012). Production of sfGFP-AID-M18BP1 cells is described below. Cells were cultured in RPMI-1640 medium supplemented with 10% fetal bovine serum, penicillin/streptomycin, and 2g/L sodium bicarbonate and maintained in a 37°C/5% CO_2_ incubator.

### Xenopus egg extracts and immunodepletions

CSF-arrested (metaphase) *Xenopus* egg extracts were prepared as described previously (Desai et al., 1999; Guse et al., 2012). In brief, eggs were washed with MMR buffer (6 mM Na-HEPES, pH 7.8, 0.1 mM EDTA, 100 mM NaCl, 2 mM KCl, 1 mM MgCl_2_, and 2 mM CaCl_2_) and then dejellied in MMR + 2% L-cysteine. Dejellied eggs were washed in CSF-XB buffer (10 mM K-HEPES, pH 7.7, 100 mM KCl, 50 mM sucrose, 2 mM MgCl_2_, 0.1 mM CaCl_2_, 5 mM EGTA), followed by CSF-XB containing 10 μg/ml LPC [leupeptin/pepstatin A/chymostatin]. Eggs were packed in a 13 × 51-mm polyallomer tube (Beckman Coulter, 326819) by a low-speed spin in a table top clinical centrifuge for 45 s. After removal of buffer, packed eggs were centrifuged in a SW55Ti rotor (Beckman Coulter) for 15 min at 10,000 rpm. The soluble cytoplasm was removed by side puncture with a 18G needle and supplemented with 50 mM sucrose, 10 μg/ml LPC, 10 μg/ml cytochalasin D, and energy mix (7.5 mM creatine phosphate, 1 mM ATP, and 1 mM MgCl_2_).

Immunodepletions of CENP-C and M18BP1 from Xenopus extract were performed essentially as described (French et al., 2017; Moree et al., 2011). Protein A Dynabeads (Invitrogen, 10001D) were washed in TBSTx (50 mM Tris-HCl, pH 7.4, 150 mM NaCl, 0.1% Triton X-100) and then coupled to affinity purified antibody for 30-60 minutes at 4°C. For 100 μL extract, 2 ug α-xCENP-C antibody (rabbit, raised and purified against xCENP-C^207-296^ (Milks et al., 2009)), or 4-5 μg α-xM18BP1 (rabbit, raised against GST-xM18BP1-2^161-415^ and purified against xM18BP1-1^161-375^ (Moree et al., 2011)) were used. An equivalent amount of whole rabbit IgG (Jackson ImmunoResearch Laboratories, Inc.) was used for mock depletions. Antibody-coupled beads were washed three times in TBSTx and then resuspended in extract. Depletions were rotated at 4°C for 1 hour. Beads were removed from extract by 2×5-minute magnetizations.

### In vitro translation and M18BP1 mutant localization

*In vitro* translated (IVT) proteins were generated in rabbit reticulocyte lysate supplemented with 20 μM methionine or 4uCi/μL [^35^S]-methionine (Perkin Elmer, NEG709A500UC) using the SP6 TNT Quick-Coupled Transcription/Translation kit (Promega, L2080) according to the manufacturer’s instructions.

To assess localization of specific M18BP1 isoforms, truncations, or mutants, 2 μL of reticulocyte lysate containing *in vitro* translated protein was added directly to M18BP1-depleted, CSF-arrested egg extract (20 μL final reaction volume). For interphase samples, reactions were supplemented with 750 μM CaCl_2_; paired metaphase reactions were supplemented with an equivalent volume of sperm dilution buffer (50 mM HEPES pH 7.7, 750 mM sucrose, 5 mM MgCl_2_, 500 mM KCl). Demembranated sperm was added to 3,000/μL and reactions were incubated at 16-18°C for 75 minutes. Reactions were then diluted in 1 mL dilution buffer (BRB-80 [80 mM K-PIPES, pH 6.8, 1 mM MgCl_2_, 1 mM EGTA], 0.5% Triton X-100, 30% glycerol) containing 150 mM KCl and incubated on ice for 5 minutes. These samples were then fixed by addition of 1 mL dilution buffer containing 4% formaldehyde and spun through cushions of 40% glycerol in BRB-80 onto poly-L-lysine-coated coverslips to be processed for immunofluorescence.

In Figure 4C to assess the effects of mitotic kinase inhibition on Myc-M18BP1-1^161-580^ localization, extract was treated with the following drugs prior to IVT protein and sperm addition: flavopiridol (10 μM; Sigma, F3055), BI2536 (10 μM; gift from Tarun Kapoor, Rockefeller University), ZM-447439 (20 μM; gift from James Ferrell, Stanford University), reversine (10 μM; AbCam, ab120921). To minimize the concentration of DMSO added to the extract, 40x drug stocks were prepared in sperm dilution buffer before addition to extract.

### CENP-A assembly assays

CENP-A assembly assays on sperm chromatin were performed as previously described (French et al., 2017; Moree et al., 2011). RNAs used to express protein in *Xenopus* egg extract were prepared with the SP6 mMessage mMachine kit (Life Technologies, AM1340). NotI linearized pCS2+ plasmids were used in the transcription reaction following manufacturer’s instructions, using double the amount of DNA. RNA was purified using RNeasy mini columns (QIAGEN). RNA was typically concentrated to at least 1 μg/μL using a SpeedVac (Savant) to avoid excessive dilution of the extract.

CENP-A assembly reactions (30 μL final volume) contained 1.5 μL xHJURP IVT protein, 0.75 μL Flag-M18BP1-1 IVT protein, and 25 ng/μL Myc-xCENP-A RNA. CSF-arrested, M18BP1-depleted extract (prepared as above) was supplemented with IVT protein and RNA and incubated at 16-18°C for 30 minutes to permit RNA translation. Cycloheximide was then added to 0.1 mg/mL to terminate translation. Reactions were then released to interphase by addition of 750 μM CaCl_2_ and supplemented with demembranated sperm chromatin to 3,000/μL. After incubation for 75 minutes at 16-18°C, assembly reactions were diluted and spun onto coverslips as described above. Quantification of CENP-A loading (centromeric Myc-CENP-A signal) was performed as described below.

### Immunoprecipitation and λ-phosphatase treatment

To assay association of full-length M18BP1-1 mutants with endogenous CENP-C, 50 μL reactions containing 5 μL *in vitro* translated Flag-M18BP1-1 in CSF-arrested or interphase M18BP1-depleted extract were added to 10 μL Protein A Dynabeads (Invitrogen) coupled to 2.5 μg mouse α-Flag antibody (Sigma).

To assay association of M18BP1-1 with Mis18α and Mis18β, 50 μL reactions containing 5 μL Flag-M18BP1-1, 2.5 μL Myc-Mis18α, and 2.5 μL Mis18β IVT proteins in CSF-arrested or interphase M18BP1-depleted extract were added to 10 μL Protein A Dynabeads (Invitrogen) coupled to 2.5 μg mouse α-Flag antibody (Sigma).

To assay association of M18BP1-1 with HJURP, 50 μL reactions containing 5 μL Flag-M18BP1-1 in CSF-arrested or interphase M18BP1-depleted extract were added to 10 μL Protein A Dynabeads (Invitrogen) coupled to 2.5 μg rabbit α-xHJURP antibody.

Following rotation for 1 hour at 4°C, beads were collected by magnetization and washed four times in TBSTx. Immunoprecipitates were eluted in sample buffer (50 mM Tris, pH 6.8, 15 mM EDTA, 1 M β-mercaptoethanol, 3.3% SDS, 10% glycerol, and 1 μg/ml Bromophenol blue), resolved by SDS-PAGE, and immunoblotted as described below.

To assess whether phosphorylation was required for the interaction between M18BP1-1 and CENP-C, precipitation reactions were prepared as above but scaled to 150 μL. 80 μL of the reaction were added to 20 μL Protein A Dynabeads (Invitrogen) coupled to 5 μg mouse α-Myc antibody (4A6; Millipore) and 40 μL was added to 10 μL Protein A Dynabeads coupled to 2.5 μg mouse whole IgG (Jackson ImmunoResearch) as a control. Immunoprecipitates were washed three times with TBSTx, then three times with LPPase buffer (50 mM Tris-HCl pH 8.0, 2 mM MnCl_2_, 0.01% Triton X-100, 10 mM NaCl). A-Myc immunoprecipitates were split to two samples, and all samples were resuspended in 50 μL LPPase buffer. Samples were then treated with 200 U lambda phosphatase (P0753S; NEB) or buffer for 30 minutes at 30°C. Beads were then washed, eluted, and processed for immunoblotting as above.

### GST-xlCENP-C^1911-1400^ and GST-hsCENP-C^723-943^ purification

GST-Prescission-xlCENP-C^1191-1400^ and GST-Prescission-hsCENP-C^723-943^ were expressed from modified pGEX plasmid (NEB) in BL21 (DE3) Codon Plus RIPL E. coli. One 2L culture was grown in 2xYT medium to OD_600_ of 0.6 at 37°C and then induced with 0.2 mM IPTG for 4 hours at 21°C. Bacteria were lysed in GST lysis buffer (10 mM sodium phosphate pH 7.4, 500 mM NaCl, 2.7 mM KCl, 1 mM EDTA, 0.5% Tween-20, 1 mM dithiothreitol (DTT), 1 mM phenylmethylsulfonyl fluoride (PMSF), 1 mM benzamidine hydrochloride, 10 μg/mL LPC [leupeptin, pepstatin A, chymostatin], 0.2 mg/mL lysozyme) by sonication and then clarified by centrifugation at 40,000 rpm in a Type 45ti rotor (Beckman) for 1 hours at 4°C. Clarified lysates were flowed over 2 mL glutathione agarose column (Sigma) equilibrated with GST lysis buffer to capture GST-tagged protein. Resin was subsequently washed with 40 column volumes lysis buffer containing 0.05% Tween-20. GST-CENP-C^1191-1400^ was then washed with 4 column volumes GST wash buffer (50 mM Tris-HCl pH 8.0, 250 mM NaCl, 1 mM DTT) and eluted with GST wash buffer containing 5 mM reduced gluathione. This protein was dialyzed against protein storage buffer (50 mM Tris-HCl pH 8.0, 250 mM NaCl, 1 mM DTT, 10% glycerol), aliquotted, and stored at −80°C.

### IVT Binding Assays

10 uL of [^35^S]-methionine-labeled IVT protein was incubated with 1.7 μM GST-xlCENP-C^1191-1400^ or GST-hsCENP-C^723-943^ in 50 mM Tris-HCl pH 8.0, 50 mM NaCl, 0.05% Triton-X 100, and 0.25 mM EDTA. The binding reaction was incubated for 1 h at 4°C. Bound material was recovered using 25 μL glutathione agarose beads, which were then washed three times in 50 mM Tris-HCl pH 8.0, 200 mM NaCl, 0.05% Triton-X 100, and 0.25 mM EDTA. Bound material was separated by SDS-PAGE and visualized by autoradiography. Gels were stained with Coomassie to visualize GST-CENP-C recovered.

Fraction bound was quantified using ImageJ (National Institutes of Health). Briefly, a rectangular region of fixed size was drawn around each band in the ‘input’ and ‘bound’ lanes and the mean integrated density measured. To estimate the local background signal, the mean integrated density was also measured for a region of the same size above the bands, and this value was subtracted from the values measured for each band. The ‘bound’ signal was divided by the ‘input’ (adjusted for the amount of the sample loaded per lane) to obtain the fraction bound.

### Histone purification

Histones were purified as described previously (Guse et al., 2012; Westhorpe et al., 2015). Briefly, histone H2A and H2B were expressed from a pET3a vector in BL21 (DE3) Codon Plus RIPL Escherichia coli (Agilent Technologies, 230280), grown in 2xYT medium (20 g/liter tryptone, 10 g/liter yeast extract, and 5 g/liter NaCl), and induced for 3 hours with 0.25 mM IPTG at an OD_600_ of 0.6. Bacteria were harvested and resuspended in lysis buffer (20 mM potassium phosphate, pH 6.8, 1M NaCl, 5 mM 2-mercaptoethanol, 1 mM PMSF, 1 mM benzamidine, 0.05% NP-40, and 0.2 mg/ml lysozyme), homogenized using an EmulsiFlex-C5 (Avestin, Inc.) followed by 1 × 30 second sonication. The lysate was centrifuged (18,000 × g, 20 min, 4°C) and the insoluble pellet was washed in lysis buffer and resuspended in unfolding buffer (7 M guanidine-HCl, 20 mM Tris-HCl, pH 7.5, and 10 mM DTT). After another round of centrifugation, the supernatant from this resuspension was dialyzed into urea buffer (6 M deionized urea, 200 mM NaCl, 10 mM Tris-HCl, pH 8, 1 mM EDTA, 5 mM 2-mercaptoethanol, and 0.1 mM PMSF). Histones were isolated by chromatography through HiTrap Q and HiTrap S columns in series. Histones were eluted from the HiTrap S column with urea buffer containing 1 M NaCl, dialyzed into water and lyophilized. To reconstitute H2A/B dimer or H3/H4 tetramer, histones were mixed and resuspended in unfolding buffer followed by dialysis into 2 M NaCl, 10 mM Tris-HCl, pH 7.6, 1 mM EDTA, and 5 mM 2-mercaptoethanol. H2A/H2B dimers and H3/H4 tetramers were then purified by size-exclusion chromatography.

Soluble CENP-A-Myc-H4 tetramer was expressed as described (Guse et al., 2012) using the pST39 vector (Tan, 2001). The initial supernatant, isolated as described above, was run through hyroxyapatite (HA) (type II 20 μM HA; Bio-Rad Laboratories) pre-equilibrated with 20 mM potassium phosphate, pH 6.8. After washing with 6 column volumes (20 mM potassium phosphate, pH 6.8, 1 M NaCl, and 5 mM 2-mercaptoethanol) tetramer was eluted with HA buffer containing 3.5 M NaCl. The elution was dialyzed into 10 mM Tris-HCl, pH 7.4, 0.75 M NaCl, 10 mM 2-mercaptoethanol, and 0.5 mM EDTA. The dialyzed protein was passed through a 1 ml HiTrap SP FastFlow column, washed, and tetramer was eluted over a 20-column volume gradient into dialysis buffer containing 2 M NaCl. Fractions containing concentrated tetramer were pooled, aliquoted, frozen in liquid nitrogen, and stored at −80°C.

### Chromatin bead reconstitution

Biotinylated 19 × 601 array DNA was purified as previously described (Guse et al., 2012). In brief, puC18 containing 19 repeats of the ‘601’ nucleosome positioning sequence (Lowary and Widom, 1998) was purified with a Gigaprep kit (QIAGEN) from SURE2 bacteria grown in Luria Broth media. The plasmid was restriction digested with EcoRI, XbaI, DraI and HindIII, and the positioning array purified by polyethylene glycol (PEG) precipitation using 0.5% incremental increases in PEG concentration, each with a 10 minute, 5,000 × g centrifugation. PEG precipitations with 19 × 601 DNA were dialyzed into TE buffer (10 mM Tris, pH 8 and 0.5 mM EDTA) and concentrated by ethanol precipitation. The EcoRI overhangs were filled with biotin-14-dATP (Invitrogen), dCTP, α-thio-dGTP, and α-thio-dTTP (ChemCyte) using Klenow fragment 3’-5’ exo-(New England Biolabs).

Reconstituted chromatin containing recombinant nucleosomes was prepared at 2 μM nucleosome concentration by salt dialysis as previously described (Guse et al., 2012). DNA and histones were pooled in high salt buffer (2 M NaCl, 10 mM Tris pH 7.5, 0.25 mM EDTA) and dialyzed over 36 hours into low salt buffer (2.5 mM NaCl, 10 mM Tris pH 7.5, 0.25 mM EDTA). H2A/H2B dimers were added at 2.2 × tetramer ratio, and 1.6 tetramers per nucleosome positioning sequence. Reconstituted chromatin was assessed by SyBr Gold (Life Technologies) staining of a 5% acrylamide native PAGE gel after chromatin digestion with AvaI.

Chromatin was bound to streptavidin-coated M-280 Dynabeads (Invitrogen, 11205D) prewashed in bead buffer (50 mM Tris pH 7.4, 75 mM NaCl, 0.25 mM EDTA, 0.05% Triton X-100 and 2.5% polyvinyl alcohol). Chromatin was bound at 2.6 fmol arrays per microgram of bead for 30-60 minutes at room temperature with constant shaking. After washing in bead buffer, beads were added to rabbit reticulocyte lysate containing *in vitro* translated protein.

### In vitro chromatin binding assays

Reticulocyte lysates containing *in vitro* translated protein were diluted with 5x CSF-XBTx (50 mM K-HEPES, pH 7.7, 500 mM KCl, 250 mM sucrose, 10 mM MgCl_2_, 0.5 mM CaCl_2_, 25 mM EGTA, 0.25% Triton X-100) to a final concentration of 1x CSF-XBTx. Protein was bound to reconstituted CENP-A or H3 chromatin by incubating 10 uL diluted IVT protein with 0.25 uL chromatin-coated beads for 60 minutes at 21°C. Beads were isolated by magnetization and washed three times with CSF-XBTx. Beads were fixed for 5 minutes at 21°C in CSF-XBTx containing 2% formaldehyde, washed into antibody dilution buffer (20 mM Tris-HCl, pH 7.4, 150 mM NaCl with 0.1% Triton X-100, and 2% bovine serum albumin), and settled on poly-L-lysine-coated coverslips for 30-60 minutes before processing for immunofluorescence microscopy.

### Human cell line generation

Transfections were carried out with FugeneHD (Promega, E2311) at a ratio of 6 μL FugeneHD:1 μg DNA according to manufacturer’s instructions.

CRISPR-Cas9 engineering was used to generate sfGFP-AID-M18BP1 DLD1s. osTIR1/TRex DLD1 cells (gift from Don Cleveland, Ludwig Cancer Institute) were transfected with px458 (Addgene 48138; (Ran et al., 2013)) modified to replace GFP with mCherry (a gift from Raj Rohatgi, Stanford University) and containing the M18BP1 sgRNA (sgRNA sequence: 5’-AAGAATTTACTTACCTCCAG-3’). These cells were cotransfected at a 1:1 mass ratio with pUC19 containing the coding sequence for sfGFP-AID flanked by sequences homologous to the 400bp upstream and downstream of the M18BP1 start codon at the endogenous locus. A silent mutation was introduced at the PAM site to prevent re-editing. At 48 hours after transfection, 25,000 mCherry-positive cells were sorted to one well of a 6-well plate by flow cytometry using a Sony SH800Z Cell Sorter to select cells expressing Cas9. Cells were expanded to a 10cm dish, and 8 days after the initial sort, single mCherry-negative and sfGFP-positive cells were resorted to each well of a 96-well plate. Successful sfGFP-AID integrants were identified by immunoblotting with a rabbit α-GFP antibody and by PCR from genomic DNA (ASON 2497: AGAATAGATCTTGTAGTTGCTGCCT; ASON 2498: GGCTGTTCTTGCTTGTCACG).

To generate complementation lines, AID-M18BP1 cells were co-transfected with pcDNA5/FRT/TO containing the full-length human M18BP1 cDNA with mRuby2 and 3xFlag fused to the C-terminus and with pOG44 encoding Flp recombinase. At 48 h after transfection, culture medium was replaced and supplemented with 50 μg/mL hygromycin to select for stable M18BP1-mRuby-Flag integrants. Hygromycin resistant colonies were obtained after 1 month of selection and pooled for localization experiments.

Degradation of endogenous CENP-C or M18BP1 was induced by supplementation of the culture medium with 1 mM indole-3-acetic acid (Fisher, 50-213-368) for 24 h.

To visualize M18BP1 degradation by immunoblotting, cells were collected by trypsinization, washed once in PBS, and then lysed by sonication in lysis buffer (25 mM Tris-HCl pH 7.4, 100 mM NaCl, 5mM MgCl_2_, 0.2% Igepal, 10% glycerol, 10 μg/mL LPC, 1 mM PMSF, 1 mM beta-glycerol-phosphate, 1 μM microcystin LR). Lysate concentrations were measured by Bradford Assay (BioRad) and normalized across samples by dilution with additional lysis buffer. After addition of sample buffer, immunoblotting was performed as described below.

For localization experiments, cells were grown on acid washed coverglass. Cells were fixed with 3.4% formaldehyde in PBS at 37°C for five minutes and then permeablized with 0.5% Triton X-100 in PBS for 5 minutes at room temperature. Coverslips were then processed for immunofluorescence.

### Immunofluorescence

Coverslips were first blocked in antibody dilution buffer (AbDil; 20 mM Tris-HCl, pH 7.4, 150 mM NaCl with 0.1% Triton X-100, and 2% bovine serum albumin) for 30 minutes. Coverslips were exposed to primary antibody diluted in AbDil for 15-30 minutes, washed three times with AbDil, and then exposed to AlexaFluor-conjugated secondary antibodies (Life Technologies; Jackson ImmunoResearch Laboratories, Inc.). When required, coverslips were blocked with 1 mg/mL whole rabbit or whole mouse IgG in AbDil followed by staining with directly-conjugated primary antibodies. Sperm nuclei and human cell coverslips were stained with 10 μg/mL Hoechst 33258 in AbDil for DNA. Coverslips were mounted in 90% glycerol, 10 mM Tris-HCl, pH 8.8, 0.5% p-phenylenediamine, sealed to a slide with clear nail polish and stored at −20°C.

Primary antibodies used were: 2 μg/mL mouse α-Flag (Sigma, F1804), 0.25-0.5 μg/mL mouse α-Myc (EMD Millipore, 4A6), 1μg/mL rabbit-α-xCENP-A, 1μg/mL rabbit-α-xCENP-C (Milks et al., 2009), 1.5 μg/ml rabbit-α-xM18BP1, 0.2 μg/mL mouse α-tubulin (Sigma, clone DM1α, T6199), and 1:100 human α-centromere serum (Antibodies, Inc., 15-234-0001). Secondary antibodies used were AlexaFluor 488-, 568-, and 647- conjugated goat α-mouse or α-rabbit (Life Technologies), and AlexaFluor 647- conjugated donkey α-rabbit (Jackson ImmunoResearch Laboratories, Inc.). All secondaries were used at 1-2 μg/mL.

### Image acquisition and processing

Imaging was performed on an IX70 Olympus microscope with a DeltaVision system (Applied Precision) a Sedat quad-pass filter set (Semrock) and monochromatic solid-state illuminators, controlled via softWoRx 4.1.0 software (Applied Precision). Images of sperm nuclei were acquired using a 60× 1.4 NA Plan Apochromat oil immersion lens (Olympus). Images of chromatin-coated beads and human cells were acquired using a 100× 1.4 NA oil immersion objective (Olympus). Images were acquired with a charge-coupled device camera (CoolSNAP HQ; Photometrics) and digitized to 12 bits. Z-sections were taken at 0.2-μm intervals (for sperm nuclei and human cells) or 0.4-μm intervals (for beads). Displayed images of sperm nuclei and human cells are maximum intensity projections of deconvolved z-stacks. Displayed images of chromatin-coated beads are a single plane from a z-stack.

Image analysis of *Xenopus* sperm (Moree et al., 2011) or chromatin-coated beads (Westhorpe et al., 2015) was performed using custom software as previously described. For sperm, at least 200 centromeres among at least 15 nuclei were counted in each condition in each experiment. For reconstituted chromatin, ~50-100 beads were quantified per condition per experiment.

### Immmunoblotting

Immunoblots were performed as previously described (French et al., 2017; Moree et al., 2011). Briefly, samples were resolved by SDS-PAGE and transferred onto polyvinylidene fluoride membrane (Bio-Rad Laboratories). For immunoblotting CENP-C, samples were transferred in CAPS transfer buffer (10 mM 3- (cyclohexylamino)-1- propanesulfonic acid, pH 11.3, 0.1% SDS, 20% methanol). All other samples were transferred in Tris-glycine transfer buffer (20 mM Tris-HCl, 200 mM glycine). Samples containing 0.75-1.5 μL IVT protein, 1.5 μL *Xenopus* egg extract, or 50 jg sonicated whole cell lysate were loaded per lane.

Membranes were blocked in 4% milk in PBSTx (10 mM sodium phosphate pH 7.4, 150 mM NaCl, 2.7 mM KCl, 0.2% Tween 20), probed with primary antibody, and detected using AlexaFluor 647-conjugated goat-α-rabbit or goat-α-mouse antibodies (Jackson ImmunoResearch Laboratories, Inc.). Detection was performed with a fluorescence imager (VersaDoc; Bio-Rad Laboratories, Inc.). In Figure 2B and 3D, M18BP1 was detected using horse radish peroxidase-conjugated goat-α-mouse secondary antibodies followed by chemiluminescence (ThermoScientific). Chemiluminscent membranes were exposed to X-ray film for 30-60s and developed with a Konica SRX-101A film processor.

Primary antibodies used were: 1 μg/mL mouse-α-Flag (Sigma, F1804), 1 μg/mL mouse-α-Myc (EMD Millipore, 4A6), 5 μg/mL rabbit-α-xM18BP1, 1 μg/mL rabbit-α-xCENP-C, 2 μg/mL rabbit-α-xHJURP (raised against GST-xHJURP^42-194^ and purified against His-xHJURP^42-194^), 2 μg/mL rabbit α-GFP (raised and purified against purified GFP), and 0.2 μg/mL mouse-α-tubulin (Dm1α; Sigma). All secondary antibodies were used at 0.4 μg/mL.

### Mass spectrometry

For identification of post-translational modifications of MBP-Prescission-xM18BP1-1^161-580^ protein was added to 100 μL M18BP1-depleted metaphase or interphase *Xenopus* egg extract at 500 nM and incubated for 75 minutes at 16-18°C. Reactions were then supplemented with 10 mM beta-glycerol-phosphate, 1 μM microcystin LR, and 1 mM benzamidine-HCl, and 80 μL of each reaction was then incubated for 2 h with 68 μg rabbit α-MBP antibody coupled to 3.4 mg Dynabeads (ThermoFisher, 14301) at 4°C. Beads were re-isolated by magnetization and washed three times with TBSTx. Proteins were eluted in sample buffer.

Isolated MBP-Prescission-xM18BP1-1^161-580^ protein separated by SDS-PAGE in 10% acrylamide, stained with Coomassie R-250 and the protein bands excised. Mass spectrometry was performed by the Taplin Mass-Spectrometry facility. Excised gel bands were cut into approximately 1 mm^3^ pieces, reduced with 1 mM dithiothreitol (DTT) for 30 minutes at 60°C and then alkylated with 5 mM iodoacetamide for 15 minutes in the dark at room temperature. Gel pieces were trypsin digested in the gel, washed and dehydrated with acetonitrile for 10 min followed by removal of acetonitrile. Gel pieces were dried in a SpeedVac and rehydrated with 50 mM ammonium bicarbonate solution containing 12.5 ng/μL modified sequencing-grade trypsin (Promega, Madison, WI) at 4°C. Samples were then incubated at 37°C overnight. Peptides were later extracted by removing the ammonium bicarbonate solution, washing once with a solution containing 50% acetonitrile and 1% formic acid, and the extracts were dried in a SpeedVac (~1 hr). The samples were then stored at 4°C until analysis. For analysis, samples were reconstituted in 5-10 μL of HPLC solvent A (2.5% acetonitrile, 0.1% formic acid). A nano-scale reverse-phase HPLC capillary column was created by packing 2.6 μm C18 spherical silica beads into a fused silica capillary (100 μm inner diameter x ~30 cm length) with a flame-drawn tip. After equilibrating the column. each sample was loaded via a Famos auto sampler (LC Packings, San Francisco CA). A gradient was formed and peptides were eluted with increasing concentrations of solvent B (97.5% acetonitrile, 0.1% formic acid).

As each peptide was eluted they were subjected to electrospray ionization before entering into an LTQ Orbitrap Velos Pro ion-trap mass spectrometer (Thermo Fisher Scientific, San Jose, CA). Eluting peptides were detected, isolated, and fragmented to produce a tandem mass spectrum of specific fragment ions for each peptide. Peptide sequences (and hence protein identity) were determined by matching protein or translated nucleotide databases with the acquired fragmentation pattern by the software program, Sequest (ThermoFinnigan, San Jose, CA). The modification of 79.9663 mass units to serine, threonine, and tyrosine was included in the database searches to determine phosphopeptides. Phosphorylation assignments were determined by the Ascore algorithm. All databases include a reversed version of all the sequences and the data was filtered to between a one and two percent peptide false discovery rate

### Protein alignment

Protein sequences homologous to the CENP-C binding domain of Xenopus M18BP1-1 were identified using NCBI PSI-BLAST with an input query of the full-length protein. Multiple sequence alignments were performed with MAFFT (version 7.305b)(Katoh et al., 2002) using default parameters.

## Acknowledgements

We thank members of the Straight lab for their critical feedback on the project and their comments on this manuscript. B.T.F. was supported by NIH T32 GM007276 and an NSF Graduate Research Fellowship. This work was supported by NIH R01 GM074728 to A.F.S.

## Author Contributions

B.T.F. performed the experiments. B.T.F. and A.F.S. designed the experiments and wrote the manuscript.

## Supplemental Figure Legends

**Supplemental Figure 1:**
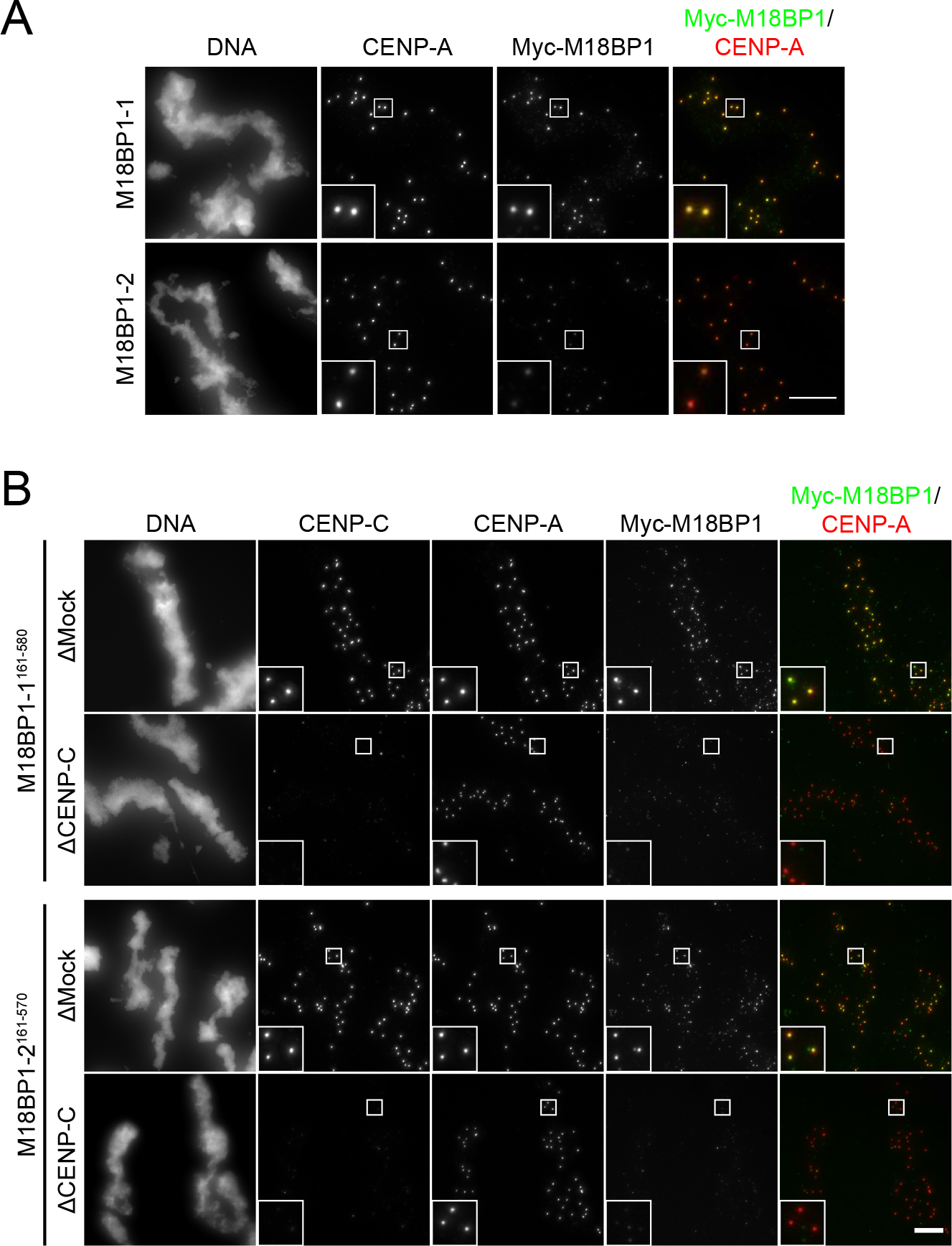
Comparison of M18BP1-1 and M18BP1-2 metaphase localization. A) Full-length M18BP1-1 and M18BP1-2 differ in their metaphase localization. Immunofluorescence images of M18BP1 isoform localization to metaphase sperm centromeres. The M18BP1 isoform is indicated to the left; immunolocalized protein is indicated above. Myc-tagged, *in vitro* translated M18BP1-1 or M18BP1-2 were incubated with sperm chromatin in metaphase-arrested Xenopus egg extract depleted of endogenous M18BP1. M18BP1-1 localizes robustly to metaphase centromeres whereas M18BP1-2 was only weakly detectable. This image represents the maximum amount of metaphase M18BP1-2 localization observed. Scale bar, 10 μm. Insets are magnified 3X. B) M18BP1-2^161-570^ exhibits CENP-C-dependent localization to metaphase centromeres similar to M18BP1-1^161-580^. Representative immunofluorescence images showing M18BP1 truncation localization (indicated at left) in metaphase extract depleted of endogenous M18BP1. In addition, extract was mock-depleted or depleted of CENP-C as indicated at left. Immunolocalized protein indicated above. Scale bar, 10 μm. Insets are magnified 3X. M18BP1-1 images are the same as those in Figure 3E.

**Supplemental Figure 2:**
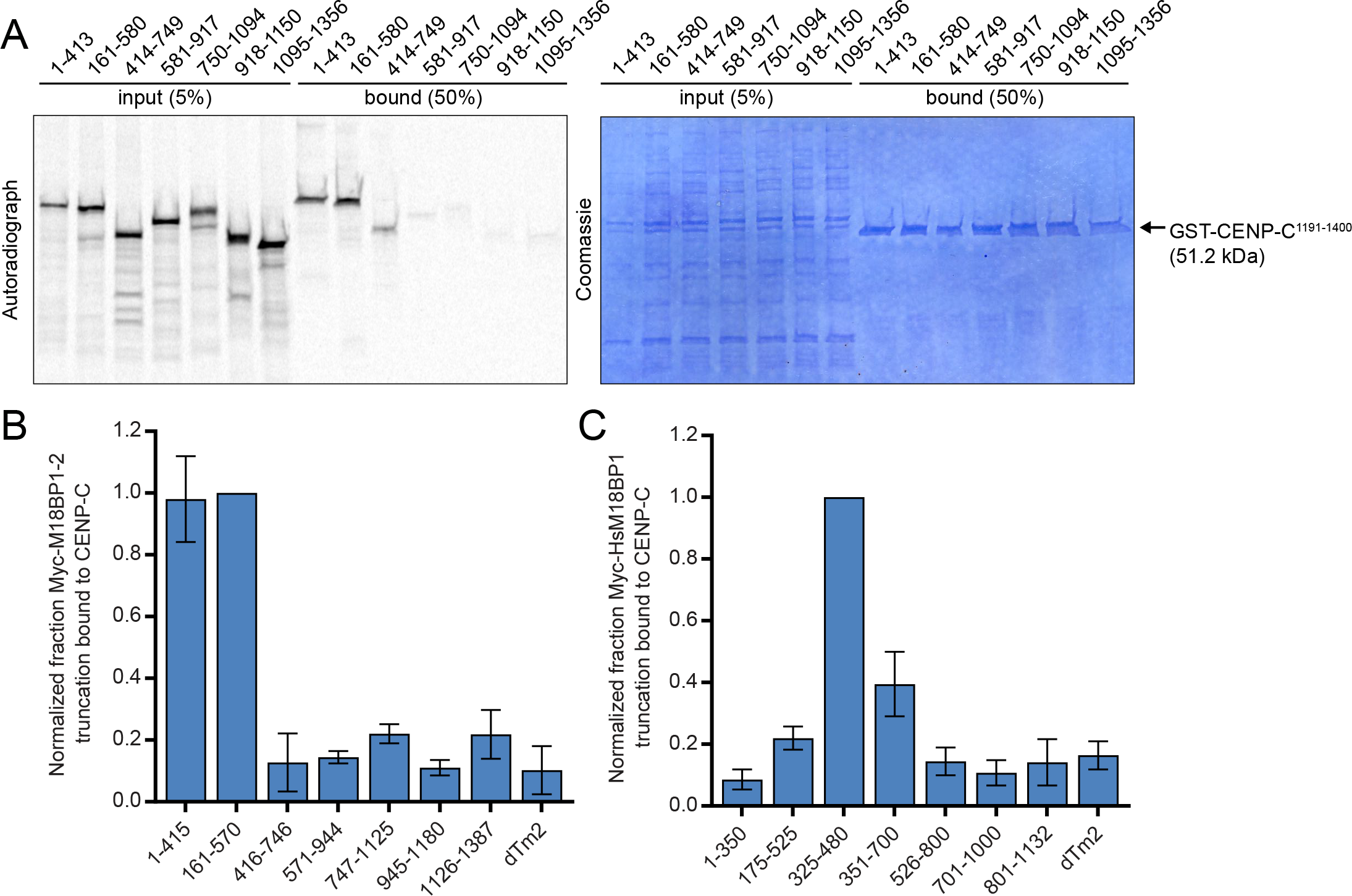
*In vitro* identification of the CENP-C binding domain in M18BP1-1, M18BP1-2, and human M18BP1. A) Reproduction of the representative GST-pulldown data from Figure 3A which includes the Coomassie stain showing equivalent recovery of GST-CENP-C^1191-1400^ for all truncations analyzed (left). B) Quantification of GST-pulldown to map the CENP-C binding domain of M18BP1-2. Myc-M18BP1-2 truncations (amino acids indicated at the bottom) were translated in reticulocyte lysate in the presence of [^35^S]-methionine and mixed with recombinant GST-CENP-C^1191-1400^. Material bound to glutathione agarose was resolved by SDS-PAGE and visualized by autoradiography to assess binding of M18BP1-2 truncations. Bound material was quantified as in Figure 3B and normalized to M18BP1-2^161-570^, the CENP-C binding domain of M18BP1-2. Notably M18BP1-2^161-415^ was not sufficient to bind CENP-C (data not shown). Error bars represent SD of two independent experiments. C) Quantification of GST-pulldown to map the CENP-C binding domain of human M18BP1. Pulldowns were performed as in (B), except radiolabeled truncations were mixed with recombinant GST-hCENP-C^723-943^, the M18BP1-binding domain on human CENP-C. Data are normalized to M18BP1^325-480^, the CENP-C binding domain on human M18BP1. Error bars represent SD of three independent experiments.

**Supplemental Figure 3:**
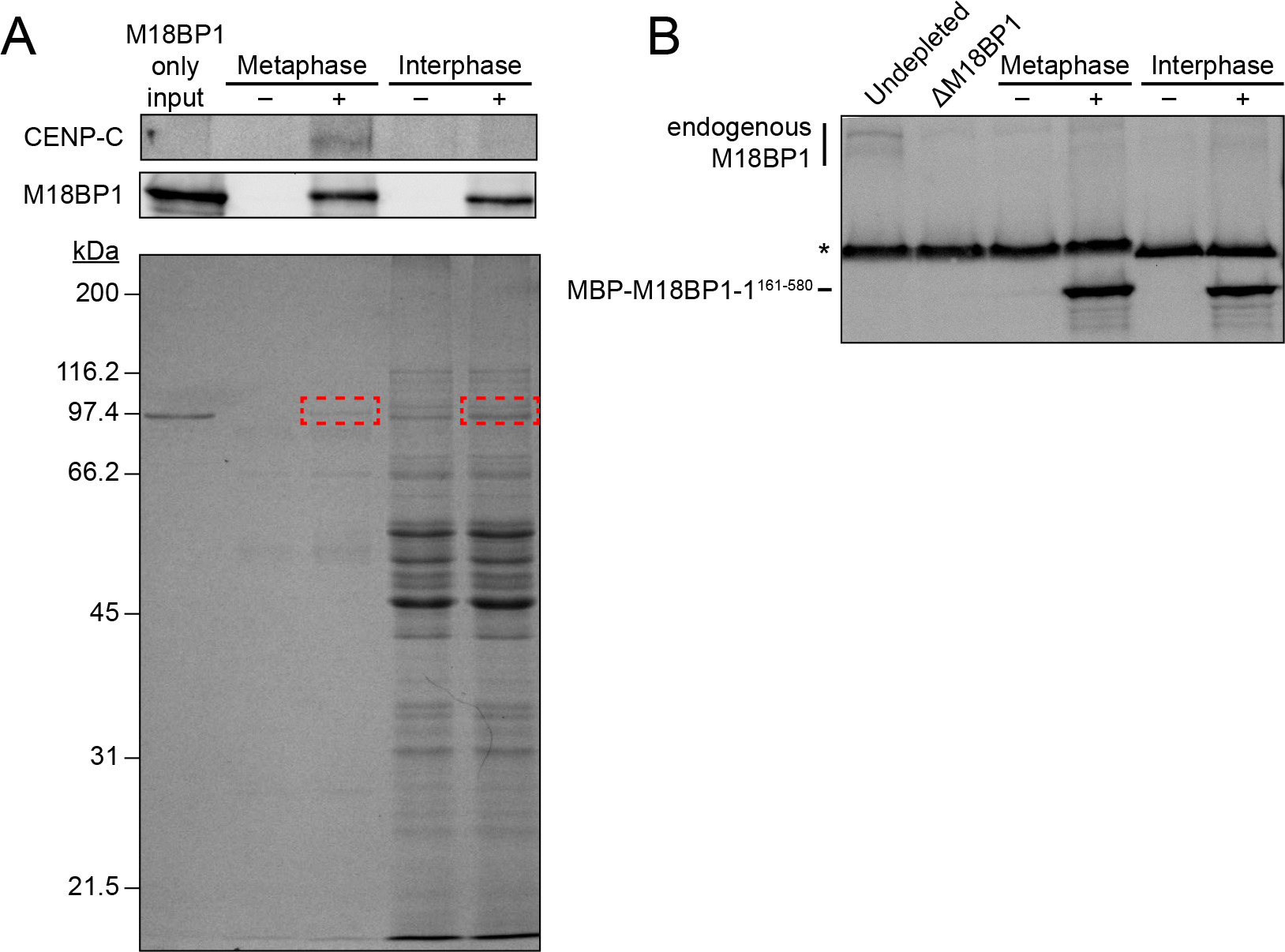
Identification of phosphosites in M18BP1-1^161-580^ by mass spectrometry. A) Pull-downs from metaphase or interphase extract depleted of M18BP1 with (+) or without (−) MBP-M18BP1-1^161-580^. (Top) Immunoblot showing that MBP-M18BP1-1^161-580^ specifically co-immunoprecipitates CENP-C from metaphase extract. (Bottom) Coomassie colloidal blue-stained gel showing material precipitated with α-MBP antibody-coated beads. Red boxes indicate bands that were excised and submitted for mass spectrometry. B) Immunoblot showing levels of MBP-M18BP1-1^161-580^ in M18BP1-depleted extract relative to endogenous M18BP1 levels (Undepleted, left lane). Nonspecific band recognized by α-M18BP1 antibody indicated by asterisk. MBP-M18BP1-1^161-580^ was ~56-fold in excess of endogenous M18BP1.

**Supplemental Figure 4:**
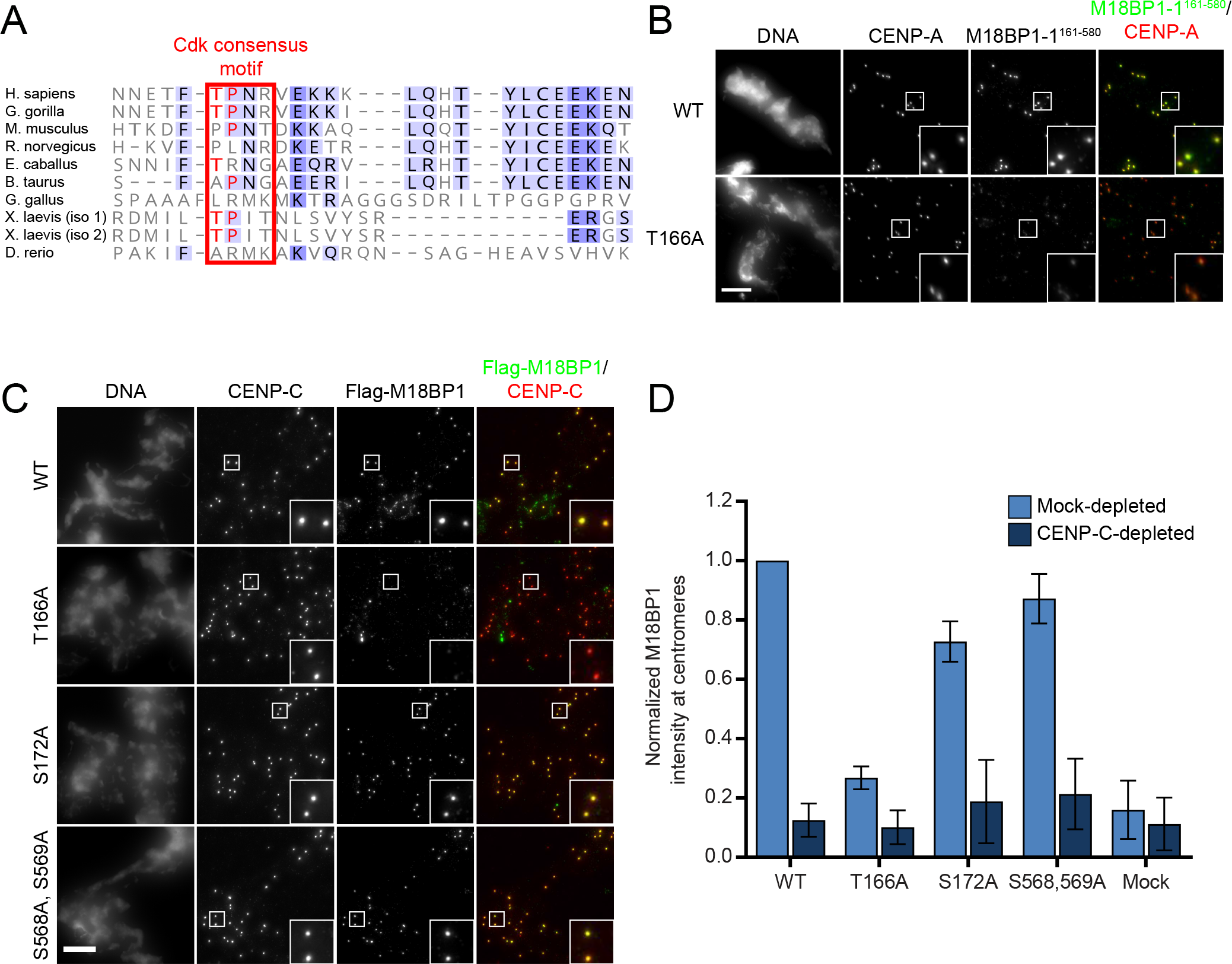
Metaphase localization of M18BP1-1 phosphomutants. A) Conservation of the *Xenopus* T166 phosphorylation site. Alignment of M18BP1 sequences among select eukaryotes. Red box indicates the position of the *Xenopus* Cdk site, with conserved T-P residues indicated in red. Darker shades of blue represent increased conservation of amino acids in alignment of ~300 M18BP1 homologues. B) Representative immunofluorescence images showing WT and T166A mutant M18BP1-1 ^161-580^ localization at metaphase sperm centromeres (see Figure 4E). Mutant species indicated at left, immunolocalized protein indicated above. Scale bar, 10 μm. Insets are magnified 3X. C) Representative immunofluorescence images showing localization of full-length M18BP1-1 phosphorylation site mutants to metaphase centromeres. M18BP1-1 species indicated at left, immunolocalized species indicated above. Scale bar, 10 μm. Insets are magnified 3X. D) Quantification of (C). Values are normalized to M18BP1-1^WT^ localization in mock-depleted extract. Error bars represent SEM of three independent experiments.

**Supplemental Figure 5:**
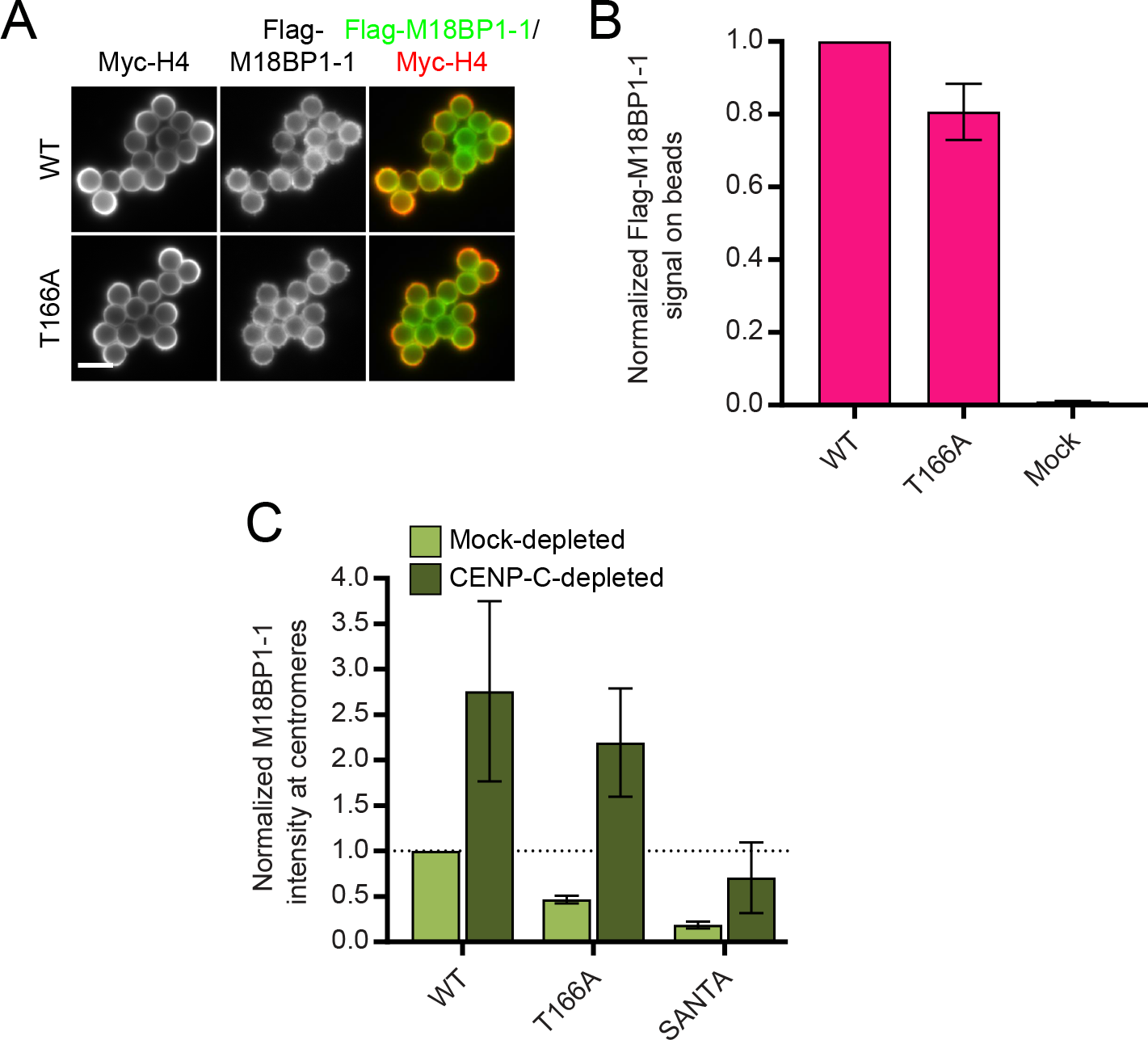
M18BP1-1^T166A^ retains the ability to bind CENP-A nucleosomes. A) Representative images of full-length Flag-M18BP1-1^WT^ or Flag-M18BP1-1^T166A^ binding to CENP-A chromatin-coated beads in reticulocyte lysate. Scale bar, 5 μm. B) Quantification of (A). Flag-M18BP1-1 fluorescence on beads was normalized to Myc-H4 signal to control for the amount of chromatin coating each bead, and then all conditions were normalized to WT. Error bars show SEM from three independent experiments. C) Metaphase targeting mutants of M18BP1 show increased localization to interphase centromeres following CENP-C depletion. French, et al. (2017) showed that this increase depends on the ability of M18BP1 to bind CENP-A nucleosomes, suggesting that the T166A and SANTA mutations do not prevent CENP-A nucleosome binding in extract. M18BP1-1 mutant species indicated below. Values are normalized to M18BP1-1^WT^ in mock-depleted interphase extract (dashed line). Error bars represent SEM of three independent experiments.

**Supplemental Figure 6:**
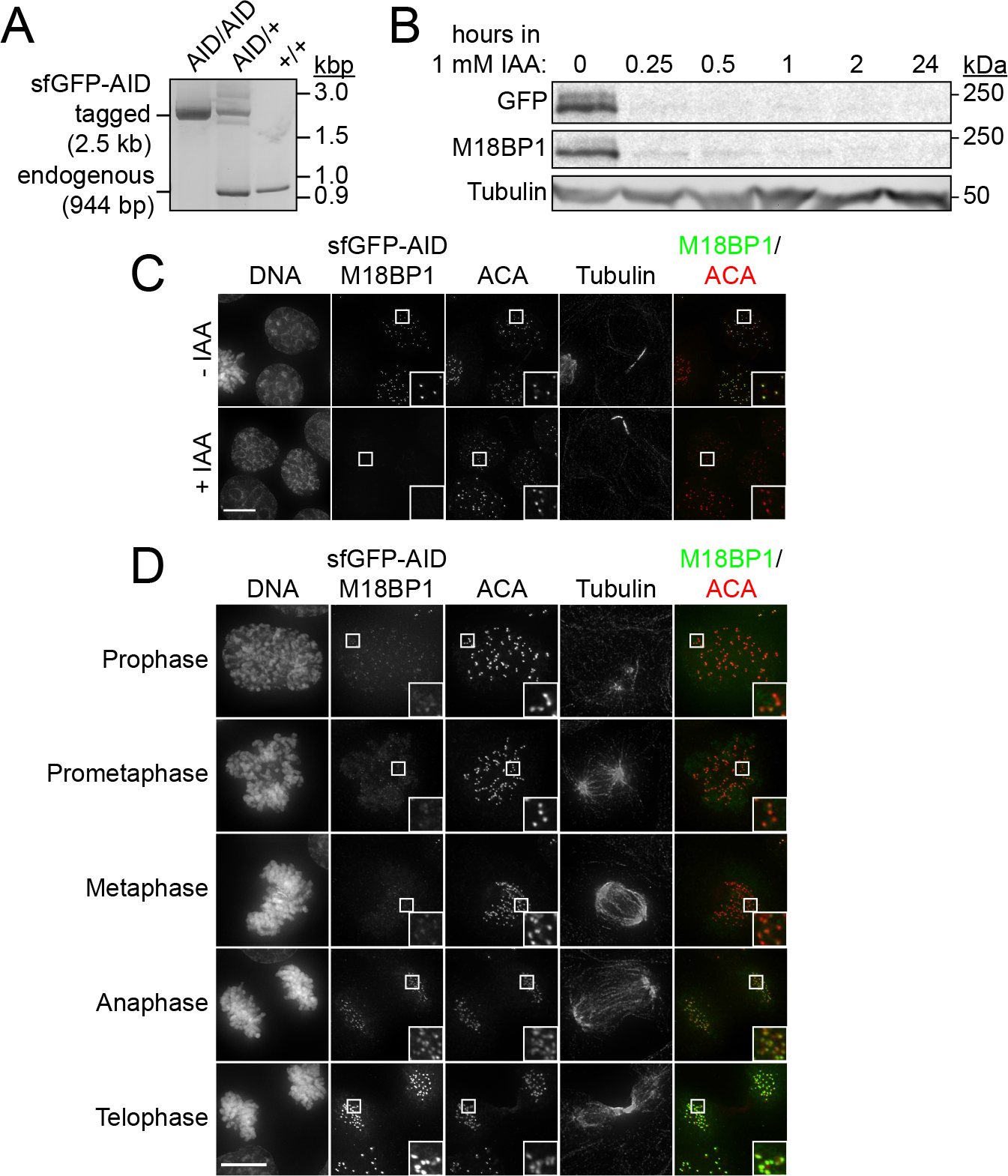
Characterization of M18BP1 localization in human cells. A) Agarose gel showing genomic PCR to assess successful integration of the sfGFP-AID tag at the endogenous M18BP1 locus. Presence of endogenous band only (944bp) indicates no integration (+/+), presence of the sfGFP-AID-M18BP1 band only (2.5 kbp) indicates successful integration at both alleles (AID/AID), and presence of both bands indicates successful integration at only one allele (AID/+). B) Immunoblot showing sfGFP-AID-M18BP1 degradation kinetics in the M18BP1^AID/AID^ cell line following supplementation of the medium with 1 mM indole-3-acetic acid (IAA). Whole-cell lysates were harvested at the indicated times following IAA addition and blotted for GFP-tagged and total M18BP1. Immunoblotted species indicated at left. C) Representative immunofluorescence images showing complete loss of G1 M18BP1 localization in the M18BP1AID/AID cell line within 1 h of IAA addition. Untreated cells are viable and show normal G1 localization of tagged M18BP1, indicating the tagged species remains functional. Midbody staining in the tubulin channel indicates early G1. ACA, α-centromere auto-antibody serum. Scale bar, 10 μm. Insets are magnified 3X. Robust centromere localization of endogenous sfGFP-AID-M18BP1 was not observed prior to anaphase/telophase in M18BP1AID/AID cells. Cell cycle state indicated at left, immunolocalized protein indicated above. ACA, α-centromere auto-antibody serum. Scale bar 10 um. Insets are magnified 3X.

## References

Barnhart, M.C., Kuich, P.H., Stellfox, M.E., Ward, J.A., Bassett, E.A., Black, B.E., and Foltz, D.R. (2011). HJURP is a CENP-A chromatin assembly factor sufficient to form a functional de novo kinetochore. J Cell Biol 194, 229–243.

Bernad, R., Sanchez, P., Rivera, T., Rodriguez-Corsino, M., Boyarchuk, E., Vassias, I., Ray-Gallet, D., Arnaoutov, A., Dasso, M., Almouzni, G., et al. (2011). Xenopus HJURP and condensin II are required for CENP-A assembly. J Cell Biol 192, 569–582.

Carroll, C.W., Milks, K.J., and Straight, A.F. (2010). Dual recognition of CENP-A nucleosomes is required for centromere assembly. J Cell Biol 189, 1143–1155.

Dambacher, S., Deng, W., Hahn, M., Sadic, D., Frohlich, J., Nuber, A., Hoischen, C., Diekmann, S., Leonhardt, H., and Schotta, G. (2012). CENP-C facilitates the recruitment of M18BP1 to centromeric chromatin. Nucleus 3, 101–110.

Desai, A., Murray, A., Mitchison, T.J., and Walczak, C.E. (1999). The use of Xenopus egg extracts to study mitotic spindle assembly and function in vitro. Methods in cell biology 61, 385–412.

Dunleavy, E.M., Roche, D., Tagami, H., Lacoste, N., Ray-Gallet, D., Nakamura, Y., Daigo, Y., Nakatani, Y., and Almouzni-Pettinotti, G. (2009). HJURP is a cell-cycle-dependent maintenance and deposition factor of CENP-A at centromeres. Cell 137, 485–497.

Fachinetti, D., Han, J.S., McMahon, M.A., Ly, P., Abdullah, A., Wong, A.J., and Cleveland, D.W. (2015). DNA Sequence-Specific Binding of CENP-B Enhances the Fidelity of Human Centromere Function. Developmental cell 33, 314–327.

Fachinetti, D., Logsdon, G.A., Abdullah, A., Selzer, E.B., Cleveland, D.W., and Black, B.E. (2017). CENP-A Modifications on Ser68 and Lys124 Are Dispensable for Establishment, Maintenance, and Long-Term Function of Human Centromeres. Developmental cell 40, 104–113.

Foltz, D.R., Jansen, L.E., Bailey, A.O., Yates, J.R., 3rd, Bassett, E.A., Wood, S., Black, B.E., and Cleveland, D.W. (2009). Centromere-specific assembly of CENP-a nucleosomes is mediated by HJURP. Cell 137, 472–484.

French, B.T., Westhorpe, F.G., Limouse, C., and Straight, A.F. (2017). Xenopus laevis M18BP1 Directly Binds Existing CENP-A Nucleosomes to Promote Centromeric Chromatin Assembly. Developmental cell 42, 190–199 e110.

Fujita, Y., Hayashi, T., Kiyomitsu, T., Toyoda, Y., Kokubu, A., Obuse, C., and Yanagida, M. (2007). Priming of centromere for CENP-A recruitment by human hMis18alpha, hMis18beta, and M18BP1. Developmental cell 12, 17–30.

Guse, A., Fuller, C.J., and Straight, A.F. (2012). A cell-free system for functional centromere and kinetochore assembly. Nature protocols 7, 1847–1869.

Hayashi, T., Fujita, Y., Iwasaki, O., Adachi, Y., Takahashi, K., and Yanagida, M. (2004). Mis16 and Mis18 are required for CENP-A loading and histone deacetylation at centromeres. Cell 118, 715–729.

Holland, A.J., Fachinetti, D., Han, J.S., and Cleveland, D.W. (2012). Inducible, reversible system for the rapid and complete degradation of proteins in mammalian cells. Proc Natl Acad Sci U S A 109, E3350–3357.

Hori, T., Shang, W.H., Hara, M., Ariyoshi, M., Arimura, Y., Fujita, R., Kurumizaka, H., and Fukagawa, T. (2017). Association of M18BP1/KNL2 with CENP-A nucleosome is essential for centromere formation in non-mammalian vertebrates. Developmental cell 42, 181–189.

Hu, H., Liu, Y., Wang, M., Fang, J., Huang, H., Yang, N., Li, Y., Wang, J., Yao, X., Shi, Y., et al. (2011). Structure of a CENP-A-histone H4 heterodimer in complex with chaperone HJURP. Genes Dev 25, 901–906.

Jansen, L.E., Black, B.E., Foltz, D.R., and Cleveland, D.W. (2007). Propagation of centromeric chromatin requires exit from mitosis. J Cell Biol 176, 795–805.

Kato, H., Jiang, J., Zhou, B.R., Rozendaal, M., Feng, H., Ghirlando, R., Xiao, T.S., Straight, A.F., and Bai, Y. (2013). A conserved mechanism for centromeric nucleosome recognition by centromere protein CENP-C. Science 340, 1110–1113.

Katoh, K., Misawa, K., Kuma, K., and Miyata, T. (2002). MAFFT: a novel method for rapid multiple sequence alignment based on fast Fourier transform. Nucleic Acids Res 30, 3059–3066.

Kim, I.S., Lee, M., Park, K.C., Jeon, Y., Park, J.H., Hwang, E.J., Jeon, T.I., Ko, S., Lee, H., Baek, S.H., et al. (2012). Roles of Mis18alpha in epigenetic regulation of centromeric chromatin and CENP-A loading. Mol Cell 46, 260–273.

Kral, L. (2015). Possible identification of CENP-C in fish and the presence of the CENP-C motif in M18BP1 of vertebrates. F1000Res 4, 474.

Lagana, A., Dorn, J.F., De Rop, V., Ladouceur, A.M., Maddox, A.S., and Maddox, P.S. (2010). A small GTPase molecular switch regulates epigenetic centromere maintenance by stabilizing newly incorporated CENP-A. Nat Cell Biol 12, 1186–1193.

Lermontova, I., Kuhlmann, M., Friedel, S., Rutten, T., Heckmann, S., Sandmann, M., Demidov, D., Schubert, V., and Schubert, I. (2013). Arabidopsis kinetochore null2 is an upstream component for centromeric histone H3 variant cenH3 deposition at centromeres. Plant Cell 25, 3389–3404.

Lowary, P.T., and Widom, J. (1998). New DNA sequence rules for high affinity binding to histone octamer and sequence-directed nucleosome positioning. Journal of molecular biology 276, 19–42.

Maddox, P.S., Hyndman, F., Monen, J., Oegema, K., and Desai, A. (2007). Functional genomics identifies a Myb domain-containing protein family required for assembly of CENP-A chromatin. J Cell Biol 176, 757–763.

McKinley, K.L., and Cheeseman, I.M. (2014). Polo-like kinase 1 licenses CENP-A deposition at centromeres. Cell 158, 397–411.

Milks, K.J., Moree, B., and Straight, A.F. (2009). Dissection of CENP-C-directed centromere and kinetochore assembly. Molecular biology of the cell 20, 4246–4255.

Moree, B., Meyer, C.B., Fuller, C.J., and Straight, A.F. (2011). CENP-C recruits M18BP1 to centromeres to promote CENP-A chromatin assembly. J Cell Biol 194, 855–871.

Muller, S., Montes de Oca, R., Lacoste, N., Dingli, F., Loew, D., and Almouzni, G. (2014). Phosphorylation and DNA binding of HJURP determine its centromeric recruitment and function in CenH3(CENP-A) loading. Cell Rep 8, 190–203.

Nardi, Isaac K., Zasadzińska, E., Stellfox, Madison E., Knippler, Christina M., and Foltz, Daniel R. (2016). Licensing of Centromeric Chromatin Assembly through the Mis18α-Mis18β Heterotetramer. Mol Cell 61, 774–787.

Ohzeki, J., Bergmann, J.H., Kouprina, N., Noskov, V.N., Nakano, M., Kimura, H., Earnshaw, W.C., Larionov, V., and Masumoto, H. (2012). Breaking the HAC Barrier: histone H3K9 acetyl/methyl balance regulates CENP-A assembly. EMBO J 31, 2391–2402.

Ohzeki, J., Shono, N., Otake, K., Martins, N.M., Kugou, K., Kimura, H., Nagase, T., Larionov, V., Earnshaw, W.C., and Masumoto, H. (2016). KAT7/HBO1/MYST2 Regulates CENP-A Chromatin Assembly by Antagonizing Suv39h1-Mediated Centromere Inactivation. Developmental cell 37, 413–427.

Pan, D., Klare, K., Petrovic, A., Take, A., Walstein, K., Singh, P., Rondelet, A., Bird, A.W., and Musacchio, A. (2017). CDK-regulated dimerization of M18BP1 on a Mis18 hexamer is necessary for CENP-A loading. Elife 6.

Perpelescu, M., Hori, T., Toyoda, A., Misu, S., Monma, N., Ikeo, K., Obuse, C., Fujiyama, A., and Fukagawa, T. (2015). HJURP is involved in the expansion of centromeric chromatin. Mol Biol Cell 26, 2742–2754.

Ran, F.A., Hsu, P.D., Wright, J., Agarwala, V., Scott, D.A., and Zhang, F. (2013). Genome engineering using the CRISPR-Cas9 system. Nature protocols 8, 2281–2308.

Sandmann, M., Talbert, P., Demidov, D., Kuhlmann, M., Rutten, T., Conrad, U., and Lermontova, I. (2017). Targeting of Arabidopsis KNL2 to Centromeres Depends on the Conserved CENPC-k Motif in Its C Terminus. Plant Cell 29, 144–155.

Session, A.M., Uno, Y., Kwon, T., Chapman, J.A., Toyoda, A., Takahashi, S., Fukui, A., Hikosaka, A., Suzuki, A., Kondo, M., et al. (2016). Genome evolution in the allotetraploid frog Xenopus laevis. Nature 538, 336–343.

Shang, W.H., Hori, T., Westhorpe, F.G., Godek, K.M., Toyoda, A., Misu, S., Monma, N., Ikeo, K., Carroll, C.W., Takami, Y., et al. (2016). Acetylation of histone H4 lysine 5 and 12 is required for CENP-A deposition into centromeres. Nat Commun 7, 13465.

Shono, N., Ohzeki, J., Otake, K., Martins, N.M., Nagase, T., Kimura, H., Larionov, V., Earnshaw, W.C., and Masumoto, H. (2015). CENP-C and CENP-I are key connecting factors for kinetochore and CENP-A assembly. J Cell Sci 128, 4572–4587.

Silva, M.C., Bodor, D.L., Stellfox, M.E., Martins, N.M., Hochegger, H., Foltz, D.R., and Jansen, L.E. (2012). Cdk activity couples epigenetic centromere inheritance to cell cycle progression. Developmental cell 22, 52–63.

Spiller, F., Medina-Pritchard, B., Abad, M.A., Wear, M.A., Molina, O., Earnshaw, W.C., and Jeyaprakash, A.A. (2017). Molecular basis for Cdk1-regulated timing of Mis18 complex assembly and CENP-A deposition. EMBO Rep 18, 894–905.

Stankovic, A., Guo, L.Y., Mata, J.F., Bodor, D.L., Cao, X.J., Bailey, A.O., Shabanowitz, J., Hunt, D.F., Garcia, B.A., Black, B.E., et al. (2016). A Dual Inhibitory Mechanism Sufficient to Maintain Cell-Cycle-Restricted CENP-A Assembly. Mol Cell 65, 231–246.

Stellfox, Madison E., Nardi, Isaac K., Knippler, Christina M., and Foltz, Daniel R. (2016). Differential Binding Partners of the Mis18α/β YIPPEE Domains Regulate Mis18 Complex Recruitment to Centromeres. Cell Reports 15, 2127–2135.

Tachiwana, H., Muller, S., Blumer, J., Klare, K., Musacchio, A., and Almouzni, G. (2015). HJURP involvement in de novo CenH3(CENP-A) and CENP-C recruitment. Cell Rep 11, 22–32.

Tan, S. (2001). A modular polycistronic expression system for overexpressing protein complexes in Escherichia coli. Protein expression and purification 21, 224–234.

Wang, J., Liu, X., Dou, Z., Chen, L., Jiang, H., Fu, C., Fu, G., Liu, D., Zhang, J., Zhu, T., et al. (2014). Mitotic regulator Mis18beta interacts with and specifies the centromeric assembly of molecular chaperone holliday junction recognition protein (HJURP). The Journal of biological chemistry 289, 8326–8336.

Westhorpe, F.G., Fuller, C.J., and Straight, A.F. (2015). A cell-free CENP-A assembly system defines the chromatin requirements for centromere maintenance. J Cell Biol 209, 789–801.

Yu, Z., Zhou, X., Wang, W., Deng, W., Fang, J., Hu, H., Wang, Z., Li, S., Cui, L., Shen, J., et al. (2015). Dynamic phosphorylation of CENP-A at Ser68 orchestrates its cell-cycle-dependent deposition at centromeres. Developmental cell 32, 68–81.

Zhang, D., Martyniuk, C.J., and Trudeau, V.L. (2006). SANTA domain: a novel conserved protein module in Eukaryota with potential involvement in chromatin regulation. Bioinformatics 22, 2459–2462.

